# Harnessing early multimodal motor training to drive motor recovery and brain-wide structural reorganization after stroke

**DOI:** 10.1101/2024.07.03.601837

**Authors:** Manuel Teichert, Sidra Gull, Karl-Heinz Herrmann, Christian Gaser, Jürgen R. Reichenbach, Anja Urbach, Christiane Frahm, Knut Holthoff, Otto W. Witte, Silvio Schmidt

## Abstract

Stroke survivors often suffer from severe motor impairments, highlighting the critical need for effective rehabilitation strategies. In experimental models, extensive task-specific motor training within the first few weeks post-stroke significantly mitigates motor deficits. However, whether more multimodal motor training approaches after stroke can restore task-specific performance in non-trained motor tasks remains largely unknown. Additionally, while stroke itself triggers structural brain reorganization, the influence of early multimodal motor training on this process remains unclear. Here, we employed T2-weighted MRI to investigate brain region-specific volumetric changes over eight weeks in rats subjected to stroke and subsequent early multimodal motor training. We found that this combination not only facilitated task-specific motor function recovery, but also induced dramatic, multi-region, brain-wide volumetric changes. Specifically, over 80 locations within 50 distinct brain regions across both hemispheres exhibited substantial volumetric alterations in a predominantly bilateral symmetric pattern. In contrast, stroke or training alone resulted in changes in 8-15 locations within 7-13 brain regions, with stroke alone primarily affecting the infarcted hemisphere. Analysis of temporal volume changes revealed two distinct trajectories in post-stroke trained rats: one of initial swelling followed by shrinkage, and another of initial shrinkage followed by swelling, suggesting an early and delayed motor learning period. Overall, our findings demonstrate that multimodal motor training early after stroke effectively restores task-specific motor function and profoundly reshapes brain structure on a brain-wide scale, offering vital insights for developing optimized rehabilitation protocols to maximize recovery in stroke patients.

## Introduction

Stroke, characterized as a sudden neurological impairment due to a disruption in cerebral blood flow, stands as one of the leading causes of mortality and disability worldwide (Langhorne et al., 2009; Stinear et al., 2020; Campos et al., 2023). It arises from the obstruction or rupture of a blood vessel, respectively, each leading to the interruption of vascular supply to brain tissue (Murphy and Corbett, 2009; Williamson et al., 2020). This causes rapid and often irreversible neuronal damage, which can manifest in lasting sensory, cognitive and motor deficits depending on the stroke’s location and severity. For instance, strokes within cortical motor regions often cause long-term contralateral motor impairments by disrupting cortical motor control (Wade, 1992; Mayo et al., 1999; Jones, 2017). Post-stroke impairments in both humans and experimental animals are signified by weakness, degraded muscle activation and other abnormalities disrupting the movement capacity and spatiotemporal coordination (Whishaw et al., 1991; Wagner et al., 2007; Klein et al., 2012; Jones, 2017). These deficits highlight the critical need for effective rehabilitation strategies to milden long-term consequences of stroke on motor function.

The majority of motor recovery or compensation (Metz et al., 2005; Jones, 2017) in both humans and rodents happens within the first three months after a stroke (Witte, 1998; Zeiler and Krakauer, 2013) before the potential for recovery or compensation progressively diminishes (Biernaskie et al., 2004; Cheatwood et al., 2008; Brown et al., 2009). Therefore, this initial period is considered as a critical window of opportunity for interventions to optimize recovery and improve outcomes (Witte, 1998; Murphy and Corbett, 2009; Krakauer et al., 2012; Stinear et al., 2020). Indeed, in experimental animals such as rodents, early and repeated task-specific motor training of the impaired limb, enriched environment that stimulates general limb use, and especially the combination of both effectively improve motor function (Biernaskie et al., 2004; Murphy and Corbett, 2009; Krakauer et al., 2012; Jeffers and Corbett, 2018; Stinear et al., 2020). However, it is less understood, whether more multimodal and holistic training approaches after stroke can also improve motor functions required in more specific and untrained tasks.

In the healthy brain, repeated motor training frequently leads to rapid volumetric increases, potentially suggestive of enhanced structural brain plasticity (Wenger et al., 2017; Li et al., 2024). Such changes have been observed in humans, for instance, following repetitive juggling training (Draganski et al., 2004), a few sessions of complex whole-body balancing (Taubert et al., 2010) or aerobic training (Erickson et al., 2011). Likewise, in rodents, volumetric increases in task specific brain regions have been found after interventions such as environmental enrichment (Scholz et al., 2015b), rotarod motor training (Scholz et al., 2015a), motor reaching (Sampaio-Baptista et al., 2013) and navigation tasks (Blumenfeld-Katzir et al., 2011).

Given that early motor training after stroke improves motor function, and that motor training *per se* often induces volumetric brain changes, it is crucial to understand whether motor training early after stroke affects brain structure as well. A deeper understanding of such effects could provide valuable insights into optimizing rehabilitation strategies. Despite this potential, these effects remain poorly understood.

In the present study, we therefore explored the effects of early multimodal motor training post -stroke on specific motor behavior and brain structure in adult rats. We found that repeated multimodal motor training, which forced general limb use, balance, sensorimotor coordination and the execution of complex motor behaviors early after a somato-motor stroke, induces a rapid recovery of task-specific, and so far untrained motor functions. In contrast, task-specific motor recovery was absent in rats subjected solely to stroke without repeated multimodal training. Strikingly, longitudinal determination of brain-region based volumetric changes within the first eight weeks post-stroke using T_2_-weighted Magnet resonance imaging (MRI) followed by Deformation Based Morphometry (DBM) revealed that motor recovery was accompanied by substantial structural reorganization across multiple brain regions located in the cortex, subcortex, and cerebellum of both hemispheres. However, in rats subjected to either stroke or training alone, structural changes were dramatically fewer and more limited in distribution. Collectively, our data demonstrate that early physical training post-stroke exerts robust effects on both motor recovery and brain structure. Moreover, our results emphasize that stroke opens a period of elevated potential for substantial structural reorganization across the whole brain, during which additional multimodal training will profoundly reshape brain structure in a brain-wide manner.

## Results

To explore the impact of early post-stroke multimodal motor training on walking behavior and brain structure, we utilized four distinct experimental groups of adult rats. In animals of the main group, we induced a stroke via photothrombosis in the right hemisphere’s somatomotor cortex. This was performed after obtaining baseline T_2_-weighted MR-images, habituating the rats to multimodal motor training on a parkour and assessing baseline skilled walking behavior using a ladder rung (LR) task, which simultaneously evaluates fore limb and hind limb function (Metz and Whishaw, 2009) (**Fig. 1 A-F**, stroke + training (“S+T”) group, n=10). After stroke, rats in this group underwent further multimodal parkour training (**Fig. 1 B, C**) three times per week for a total of eight weeks, during which they were MR-imaged periodically and tested on the LR task (**Fig. 1 A**). The parkour for multimodal motor training was designed to force general limb use, balance, sensorimotor coordination and the execution of complex motor behaviors (**Fig. 1 B**). In addition, starting from week two we gradually increased the difficulty of the parkour in order to keep the demand to finish the parkour at a high level. Thus, our parkour training combined several therapeutic components with proven benefits in stroke recovery such as repetitive, temporally spaced and variable motor practice, multisensory stimulation (visual, somatosensation) and increased difficulty (Maier et al., 2019). To isolate the effects of stroke and training alone, we established two additional groups: One group, which was only subjected to stroke alone (’S only’ group, n=16), but not parkour training and another group solely subjected to multimodal parkour training only (’T only’ group, n=10). The latter group received a sham photothrombosis and rats of both groups underwent the LR task. Additionally, a control group (‘Control’, n=8) was only subjected to T_2_-weighted imaging, received a sham phototrombosis but no training, and its performance was evaluated in the LR task (**Fig. 1 A**).

**Figure 1:**
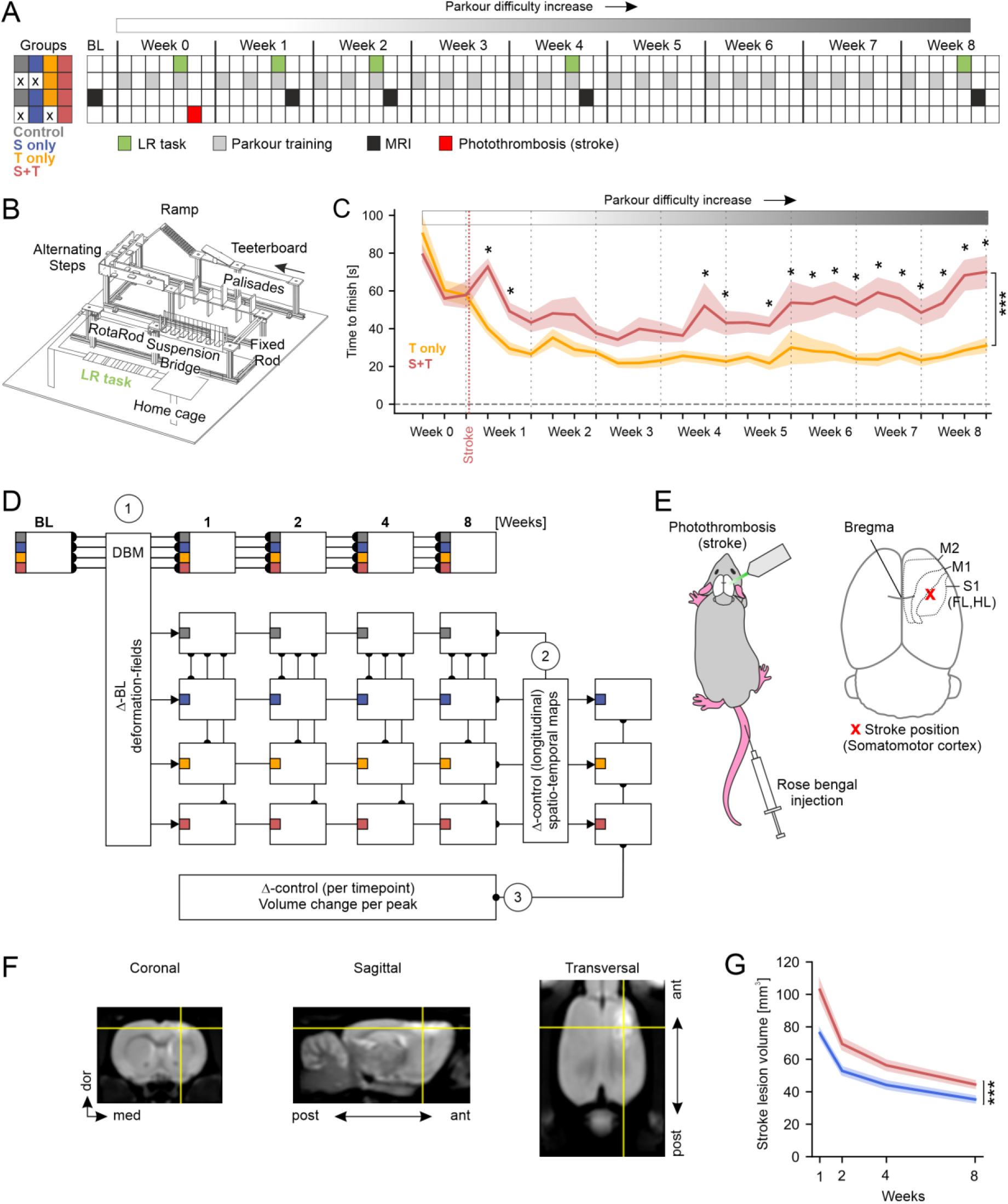
Study design. (**A**) Left: Color code of the different control and experimental groups. Filled squares indicate the presence and crosses indicate the absence of a specific treatment throughout the eight weeks study period. Right: Timeline depicting the time points of specific treatments throughout the eight weeks study period. (**B**) Schematic of the parkour utilized for physical training. Note that the LR task at the end of the parkour was only installed when walking behavior was assessed. Otherwise, a normal jetty was inserted in its place. Black arrow indicates starting position and directionality of the parkour. (**C**) Line graph showing the average time required to complete the parkour over time (± s.e.m.). Note, that only rats of the “T only” and “S+T group” participated in this training. The time to finish was significantly different within (p<0.05) and between the groups (p<0.001, two way repeated measure ANOVA with Tukey HSD). Post-hoc Tukey HSD test revealed significant differences between the groups at specific time-points (p<0.05) (**D**) Schematic of the steps involved in MRI data processing and subsequent statistical analysis. 1) Deformation Based Morphometry (DBM): brains scanned at each time-point were morphed to their own baseline. 2) Detection of spatio-temporal interactions between the control (‘Control’) and interventional groups (‘S only’, ‘T only’ and ‘S+T’). 3) Detection of the direction of volumetric changes (swelling or shrinkage) by correcting data of interventional groups to the control group. (**E**) Schematic of the procedure for stoke induction via photothrombosis, targeting the somatomotor cortex based on stereotaxic coordinates (Paxinos and Watson, 2005). The red cross indicates the position of stroke induction relative to Bregma (see Methods). M2, secondary motor cortex; M1, primary motor cortex; S1 (FL, HL), primary somatosensory cortex of the fore- and hind limb. (**F**) Representative T2-weighted MRI images of a rat brain eight weeks post-stroke. The intersections of the yellow lines mark the focal point of the stroke’s impact. (**G**) Line plot showing the average stroke volume over time as estimated by the Cavalieri method (see Methods). Volumetric changes over time were significant within and between the groups (p<0.001, two way repeated measure ANOVA with Tukey HSD). *, p<0.05; ***, p<0.001.

After baseline imaging, rats from all groups underwent further T_2_-weighted MRI at one, two, four, and eight weeks post-stroke or sham procedure. To analyze changes in brain volume over this time period, imaged brains were morphed to match their own baseline images using DBM (**Fig. 1 D**). Moreover, to associate structural changes with specific brain regions, all brain images were registered to the Paxinos rat brain atlas (Paxinos and Watson, 2005; Gaser et al., 2012)(see Methods). Subsequently, morphed brains from each group were compared with those of the control group at each time point to identify significant alterations in brain volume (**Fig. 1 D**, see Methods).

During repeated multimodal parkour training, we consistently measured the time required to complete the parkour for both post-stroke trained rats and rats that solely underwent multimodal parkour training (**Fig. 1 B, C**). Notably, rats of both groups remained motivated, consistently navigating the entire parkour without significant pauses. Although post-stoke trained rats displayed noticeable walking impairments (visible in the left fore and hind paw) and took significantly longer to complete the parkour compared to rats without a stroke, all reached the end (home cage) in each trial (**Fig. 1 C**). In detail, post-stroke trained rats exhibited a significant increase in the time required to complete the parkour immediately after the stroke which was followed by a gradual decrease until week three. From that point, the required time gradually increased again until week eight, coinciding with the increasing difficulty of the parkour. In contrast, rats subjected solely to training initially showed a decrease in the time to finish the parkour until weeks two to three, after which their completion time remained stable for the rest of the study period (**Fig. 1 C**). In rats trained post-stroke, the initial increase in time immediately after the stroke, followed by a gradual reduction until week three, suggests that rats in this group needed to relearn how to navigate the parkour due to their motor impairments. The subsequent stepwise increase in time indicates difficulties in adapting to progressively more challenging physical tasks achieved by increasing the parkour’s difficulty. Collectively, these observations suggest that post-stroke trained rats experienced two distinct periods of motor learning and adaptation: an early phase of relearning basic motor skills and a later phase of adapting to increased task complexity.

Finally, our comparison of the stroke lesion volumes in T_2_-weighted MRI images revealed an initial increase in post-stroke trained rats compared to rats that did not perform parkour training after stroke. However, over the assessed time course, in both groups stroke volume gradually decreased (**Fig. 1 F, G**). Both training induced increase of stroke volume as well as its gradual reduction over time have been repeatedly observed, and are presumably attributable to a to a high vulnerability of the region surrounding the injury to behavioral pressure (Kozlowski et al., 1996; Risedal et al., 1999) and to healing processes such as the formation of a glial scar surrounding the infarction (Jolkkonen et al., 2007; Shimada et al., 2011; Li et al., 2014).

### Multimodal motor training early after stroke improves task-specific motor behavior

Next, we explored the impact of early post-stroke multimodal motor training on task-specific motor performance using the LR task (**Fig. 2 A**). Specifically, we assessed the precision of all four paws’ placement as the rats navigated a horizontal ladder (**Fig. 2 B**). Control rats, that neither received a stroke nor were trained on the parkour, consistently demonstrated high precision and experienced no difficulties in paw placement at any time point (**Fig. S1**). Similar observations were made in rats that solely underwent parkour training, with only marginal changes in paw placement performance compared to control rats (**Fig. 2 C-F**). In contrast, one week after stroke, both rats that only received a stroke and post-stroke trained rats exhibited a significant decline in paw placement performance, particularly affecting the left fore- and hind paws, with the most pronounced decline observed in the left hind paw (**Fig. 2 C, E**). However, while the performance with these paws remained impaired in rats that did not undergo multimodal motor training after stroke, the performance significantly recovered to control levels in post-stroke trained rats (**Fig 2 C, E**). The performance with the right fore and hind paw largely remained unchanged after stroke and showed only minor but significant differences between the groups over time (**Fig. 2 D, F**). Collectively, these results imply that multimodal motor training early after stroke recovers motor execution initially impaired after stroke in a task specific manner. This is further suggestive of a transfer of improved motor function obtained during multimodal motor training to a more specific motor task.

**Figure 2:**
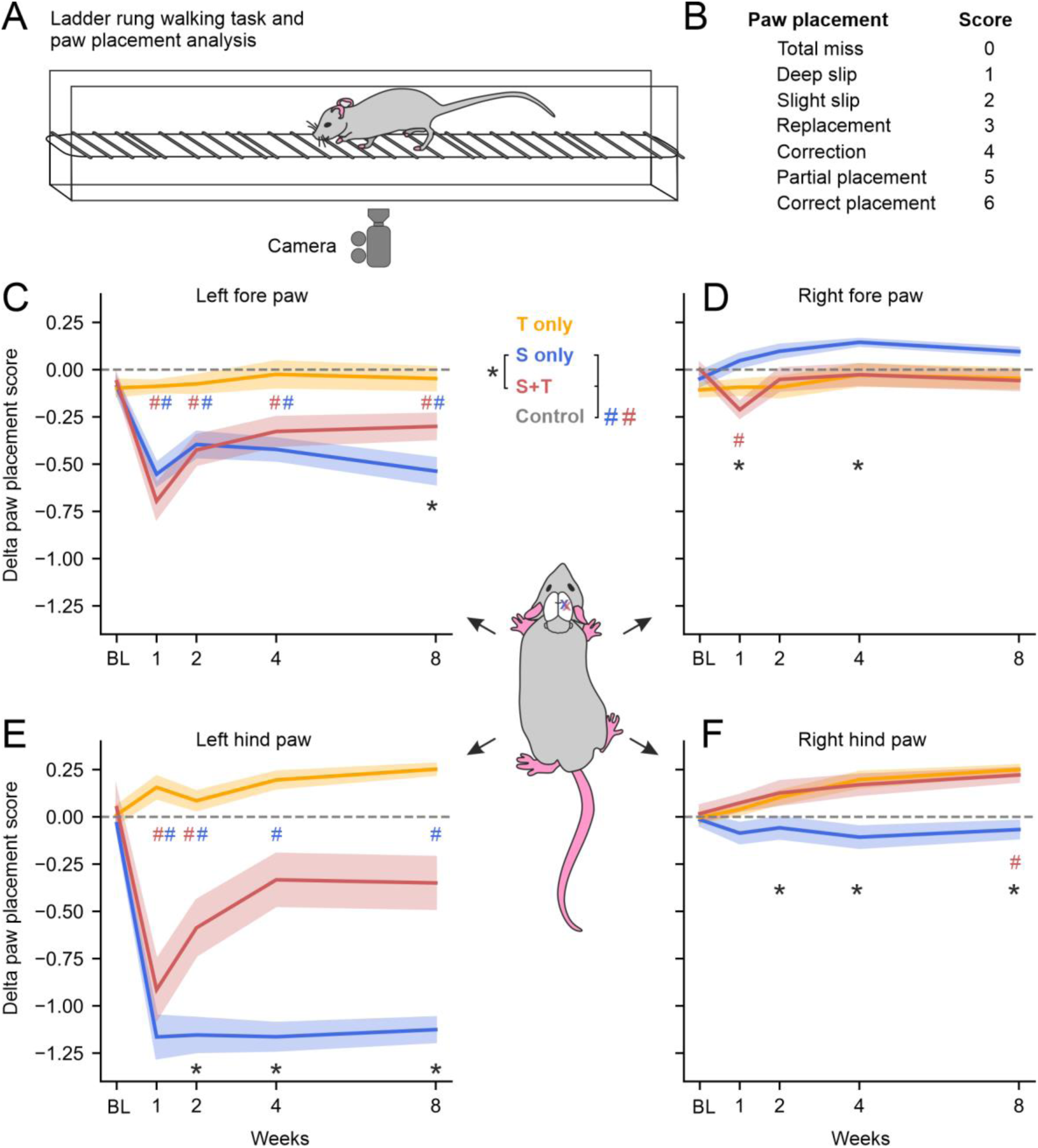
Multimodal training early after stroke induces rapid recovery of motor behavior. (**A**) Schematic of the ladder rung task. The animal’s walking behavior was recorded by a camera from below for subsequent analysis. (**B**) Paw placement scores used for quantification of the precision in placing all different paws on the ladder. (**C**-**F**) Line plots showing the differences in the paw placement score of each individual paw over time compared to control rats. Plotted is the mean ± s.e.m. of each group. Crosses in the right hemisphere of the rat’s brain indicate stroke location. For statistical analysis, we used a two way repeated measure ANOVA with Tukey HSD. *, p < 0.05 (‘S only’ vs ‘S+T’); #, p<0.05 (‘Control’ vs. ‘S only’ or ‘S+T’).

### Multimodal training early after stroke is accompanied by brain-wide structural reorganization

It is well established that motor training causes behavioral improvements that often correlate with changes in the volume of functionally involved brain regions (Wenger et al., 2017). Thus, we next investigated whether behavioral improvements in post-stroke trained rats are accompanied by structural brain changes as well. For this, we first searched for brain-region specific structural changes in the interventional groups that showed significant volumetric differences over the whole study time compared to control rats and independent of the direction of volume changes (swelling or shrinkage, see Methods).

In rats that only received a stroke, we detected significant structural changes in 7 cortical and 4 subcortical brain regions. In one of the latter, the caudate putamen (Cpu), structural alterations occurred bilaterally. In addition, volumetric changes were also detected within the white matter (**Fig. 3 A, C**). In rats that exclusively underwent multimodal parkour training, we detected 5 cortical and two cerebellar regions with significant structural alterations, while in one cortical and two cerebellar regions, namely the paraflocculus (PFI) and the Crus of the ansiform lobule (Crus), volume changes appeared to be bilateral (**Fig. 3 A, C**). These data indicate that both, stroke and training alone cause changes in brain volume, although, primarily in different sets of brain regions.

**Figure 3:**
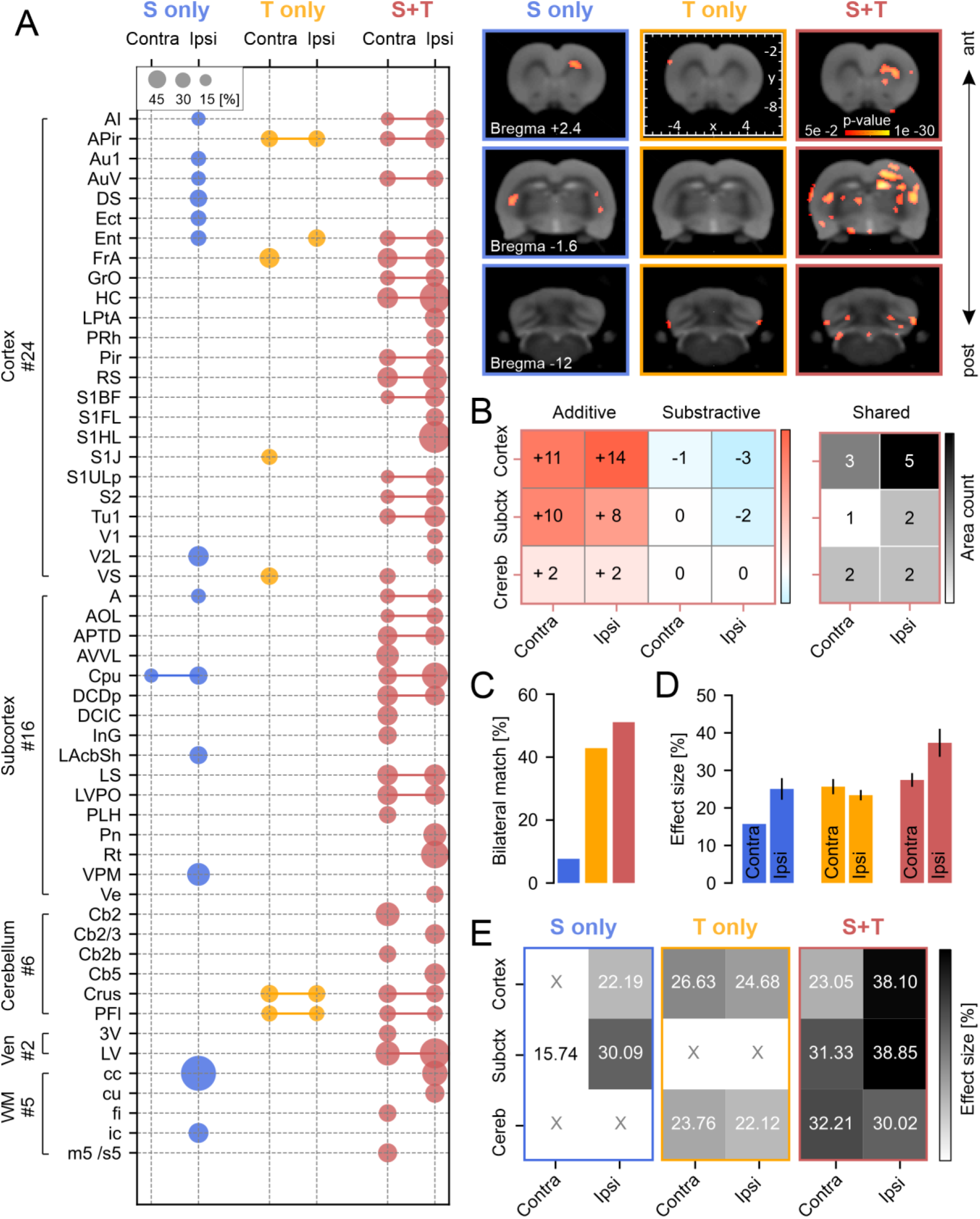
Multimodal training early after stroke induces brain-wide structural reorganization within multiple brain areas. (**A**) Left: Brain region-based analysis of the effect size of significant structural changes. The size of each dot indicates the effect size within a specific brain region and connecting lines indicate the appearance of bilateral effects. Right: Representative statistical parametric maps showing significantly affected brain regions superimposed with coronal sections of the atlas-based (Paxinos and Watson, 2005) reference brain ordered along the anterior-posterior brain axis (T2-weighted MRI images). (**B**) Heat maps showing the number of brain regions with significant structural alterations only detected in *’*S+T*’* rats, but neither in *’*S only*’* or *’*T only*’* rats (additive, color-coded in red), the number of brain regions in which significant structural changes disappeared due to the combination of stroke and training (*’*S+T*’*, subtractive, color-coded in blue), and the number of brain regions with significant structural alterations present in *’*S+T*’* rats and/or rats of the *’*S only*’* or *’*T only*’* group (shared, color coded in black). (**C**) Bar plot showing the percentage of brain areas with significant structural changes in both hemispheres on all different areas with significant structural changes. (**D**) Bar plot showing the average effect size in both brain hemispheres (± s.e.m). (**E**) Heat map showing the average effect size in cortical, subcortical and cerebellar brain regions. The number given in each square indicates the average effect size. Statistical analysis is described in the Methods (DBM). For abbreviations see **Supplementary Table 1**.

Strikingly, rats trained post-stroke exhibited significant volume changes in 20 cortical, 14 subcortical and 6 distinct cerebellar regions. Additionally, significant changes were also visible in the ventricle and the white matter (**Fig. 3 A**). Importantly, in more than 50% of the regions structural changes were detected bilaterally (**Fig. 3 C**). These data indicate that multimodal motor training early after stroke causes significant structural changes in multiple, widely distributed brain regions across both hemispheres, indicative of a synergistic effect of stroke and subsequent training on brain volume dynamics.

In our further analysis, we focused on cortical, subcortical, and cerebellar regions, where most of the significant structural changes were observed (**Fig. 3 A**). We identified numerous brain regions showing significant structural changes exclusively in post stroke trained rats (**Fig. 3 B**, additive). Conversely, some brain regions only showed significant volume changes in rats exclusively subjected to stroke or training, but lacked changes in post-stroke trained rats (**Fig. 3 B**, subtractive). Additionally, we found that some brain regions exhibited significant volume changes in rats of all groups (**Fig. 3 B**, shared). These data imply that early post-stroke motor training mainly triggers unique structural reorganization, but also leads to a loss or maintenance of volumetric changes in brain regions that only occurred after stroke or training alone.

Significant volumetric changes in the detected brain regions showed both volumetric increases (swellings) and decreased (shrinkages). In order to get insights into the maximal range of structural alterations, we calculated the total percent volume change in all brain regions over time, which we here refer to as the ‘effect size’ (**Fig. 3 A, D**). Comparison within the three groups revealed that in both, rats subjected to stroke alone and rats trained post-stroke, the average effect size was lower in the contralateral compared to the ipsilateral hemisphere, while in rats subjected to training alone no obvious hemispheric differences were detected (**Fig. 3 D**). Further quantification revealed that in post-stroke trained rats the average effect size was strongly pronounced primarily in ipsilateral cortical, subcortical and cerebellar regions compared to rats of the other groups (**Fig. 3 E**). Together, our data suggest that early post-stroke multimodal training intensifies structural adaptions, producing pronounced region-specific volume changes.

### Multimodal training post-stroke causes volume changes in multiple sub-locations within brain regions

So far, we have demonstrated that post-stroke motor training induces substantial volume changes in various cortical, subcortical, and cerebellar regions. However, our analysis also revealed that such changes often occurred at multiple sub-locations within individual regions, which we here refer to as individual ‘peaks’. Consequently, we analyzed the spatial distribution and effect size of such peaks. In rats subjected to stroke alone, these peaks predominantly appeared in posterior cortical and subcortical locations of the ipsilateral hemisphere, but not in the cerebellum, whereas in rats subjected solely to multimodal training, such peaks occurred in both anterior and posterior cortical locations, as well as in the cerebellum, but were absent in subcortical areas (**Fig. 4 A, B**). In contrast, in rats trained post-stroke, peaks were distributed brain-wide, with predominant occurrences in the cortex, followed by the subcortex and cerebellum across both hemispheres (**Fig. 4 A, B**). Importantly, in all these nodes within both hemispheres, the number of peaks detected in post-stroke trained rats was consistently higher compared to those found in rats of the other two groups (**Fig. 4 A, B**). Further analysis also revealed nuanced effects of post-stroke training regarding the peak counts within specific brain regions, compared to rats of the other groups. For instance, within regions such as the piriform cortex (Pir), the barrel field and the upper limb region of the primary somatosensory cortex (S1BF and S1ULp, respectively), all only found in rats trained post-stroke, we detected more than one and up to 5 individual peaks (**Fig. 4 C**). In contrast, in rats solely subjected to stroke or training, most regions contained only one peak. Additionally, in brain regions that showed significant structural changes in post-stroke trained rats, but also in rats of at least one of the other groups (entorhinal cortex (Ent), Cpu, Crus and PFI), the number of peaks was typically higher in post-stroke trained rats that compared to rats of the other groups (**Fig. 4 C**). In sum, our data suggest that compared to stroke or motor training alone, their combination significantly enhances structural reorganization, leading to more widespread and numerous volumetric changes across and within affected brain regions.

**Figure 4:**
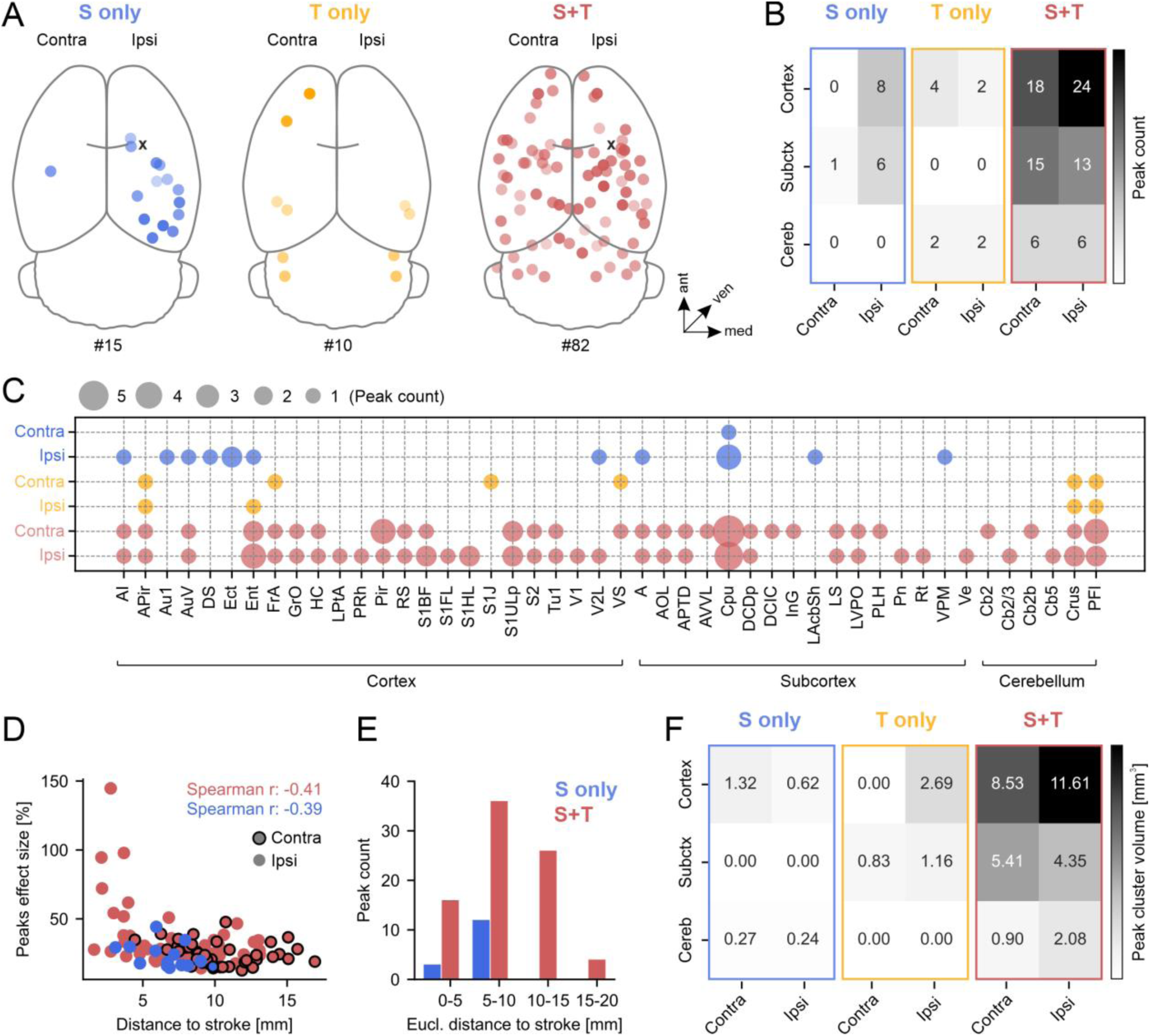
Multimodal training post-stroke induces widespread volumetric changes that encompass greater brain volume. (**A**) Three dimensional plots showing the distribution of significant peaks across the brain. Darker points indicate more dorsally and paler data pints indicate more ventrally peak locations. Black crosses indicate stroke position. The numbers below the depicted brains indicates the total peak count. (**B**) Heat map displaying the peak count in the cortex, the subcortex and the cerebellum. Numbers within the squares indicate peak count. (**C**) Dot plot showing brain regions with significant volume changes over the time studied. The dot size indicates the number of detected peaks within each brain area. For abbreviations see **Supplementary Table 1**. (**D**). Scatterplot showing the effect size as a function of the Euclidean distance to the stroke center. Values for r indicate the Spearman’s rank correlation coefficient. (**E**) Bar plot showing the peak count at different distances from the stroke. (**F**) Heat map showing the peak cluster volume in the cortex, the subcortex and the cerebellum. Statistical analysis is described in the Methods (DBM).

Next, we asked whether the effect size of each individual peaks is dependent on the Euclidean distance of each peak from the center of the stroke. Indeed, in both groups—rats subjected to stroke alone and post-stroke trained rats—we observed a negative correlation, with peaks closer to the stroke center displaying higher effect sizes and those further away exhibiting lower effect sizes (**Fig. 4 D**). Importantly, the highest peak effect sizes occurred within a 5 mm radius around the stroke center. These changes may be attributable to the damage of neuronal tissue, the observed motor training induced alterations of stroke volume, or local edema in its direct proximity (Slivka et al., 1995; Witte et al., 2000; Koch et al., 2019). However, the effect sizes of peaks between 5-20 mm away from the stroke center appeared to be of similar size. These changes are likely not influenced by neuronal damage or the stroke *per se* but rather reflect structural alterations of healthy brain tissue. Moreover, in both groups, the number of peaks gradually increased from 0 to 10 mm away from the stroke center, followed by a gradual decrease between 10-20 mm around the stroke (**Fig. 4 E**), suggesting that the number of local volumetric alterations is dependent on their distance to the stroke center.

Each individual location that showed volumetric changes was composed of multiple, seamlessly connected voxels, here termed ‘peak clusters’. Consequently, we calculated the volume of each peak cluster within the brain across the three groups. Our analysis revealed that, in rats trained post-stroke, the mean volume of these peak clusters was substantially larger across all brain nodes—the cortex, subcortex, and cerebellum in both hemispheres— compared to those in rats subjected only to stroke or training (**Fig. 4 F**). These findings indicate that early post-stroke motor training mobilizes larger volumes of brain tissue for structural reorganization.

### Initial structural volume changes invert their directionality over time

As previously noted, brain volume can increase (swell) or decrease (shrink) following specific interventions (Wenger et al., 2017). Consequently, we investigated the directionality of volume changes at one, two, four, and eight weeks to determine how stroke, motor training, and their combination dynamically influence brain volume over time.

In rats subjected solely to stroke, the majority of peaks exhibited significant volume swellings after one week, with only one showing significant shrinkage (**Fig. 5 A, B**). By two weeks, most changes lost significance, but regained significance between four and eight weeks. Interestingly, all peaks that initially showed significant swellings reversed their direction of volume change between one and four weeks post-stroke. The one peak that initially exhibited significant shrinkage also reversed its volumetric change (**Fig. 5 C**, **Fig. S2 A**). In contrast, rats that exclusively underwent multimodal training exhibited minor and non-significant volume changes, including both shrinkages and swellings, after the first week. Over the subsequent weeks, changes were primarily directed towards significant shrinkage, with only a few exhibiting significant swellings (**Fig. 5 A-C**, **Fig. S2 B**).

**Figure 5:**
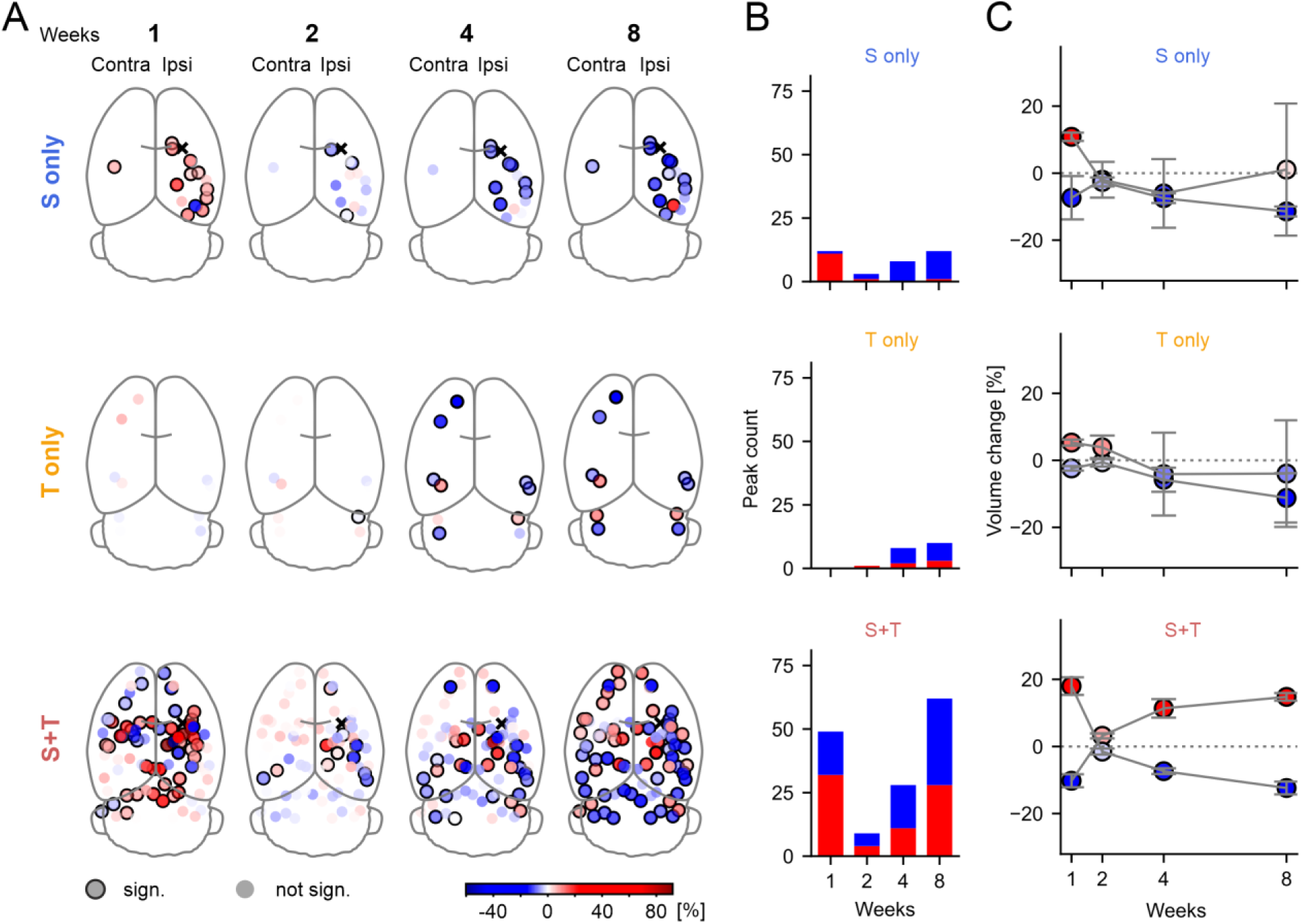
Initial structural volume changes invert their directionality within the first weeks after stroke. (**A**) Three dimensional plots showing volume changes of detected peaks (red, swelling; blue, shrinkage) over time. Data points surrounded by a black circle display significant volumetric changes. Black crosses indicate stroke position. (**B**) Bar plots showing the count of significant volume changes over time. (**C**) Line plots showing the average volume changes over time ± s.e.m. Data were selected based on the direction of volume changes (either shrinkage or swelling, <0 or >0) at week one. Statistical analysis is described in the Methods section (DBM). Values of percentage volume changes for each peak are provided in **Supplementary Table 2-4**.

In rats trained post-stroke, we observed numerous substantial volume changes within one week, with comparable numbers of peaks exhibiting initial shrinkage or swelling (**Fig. 5 A, B**). Similar to the stroke-only group, these volume changes largely became insignificant by the second week but regained significance from the fourth to the eighth week post-stroke (**Fig. 5 A**). Remarkably, out of 81 peaks identified, 80 reversed their direction of volume change between the first and fourth week: nearly all peaks initially showing shrinkage transitioned to swelling, and vice versa (**Fig. 5 C**, **Fig. S2 C**). The most pronounced alterations in volume magnitude occurred between the first and fourth week, while changes were minor from the fourth to the eighth week post-stroke (**Fig. 5 C**, **Fig. S1 C**). In summary, these results suggest that the directionality and magnitude of brain volume changes in rats are dynamically influenced by the type of intervention, with the most clearly pronounced pattern revealed in post-stroke trained rats. In addition, given that motor training often causes swellings in specific brain regions (Wenger et al., 2017), the observed two temporally distinct swelling periods argue for two motor learning periods in post-stroke trained rats, one initial and one delayed, while in rats exclusively subjected to stroke only one initial learning period may take place early after stroke.

### Early multimodal training post-stroke induces two distinct temporal trajectories of brain volume swellings and shrinkages over time

Having demonstrated that initial volumetric changes—swellings and shrinkages—typically reverse over time, we further investigated how these distinct temporal trajectories are distributed across the brain and within specific regions among the three groups. To delineate these temporal patterns, we first conducted a k-means cluster analysis of the pooled volumetric data of all groups. Despite the different temporal dynamics revealed by the previous analysis (see Fig. 5), cluster analysis indicated the presence of two dominant temporal trajectories in all groups: one characterized by initial swelling followed by gradual shrinkage, and another by initial shrinkage followed by gradual swelling (**Fig. 6 A, B**).

**Figure 6:**
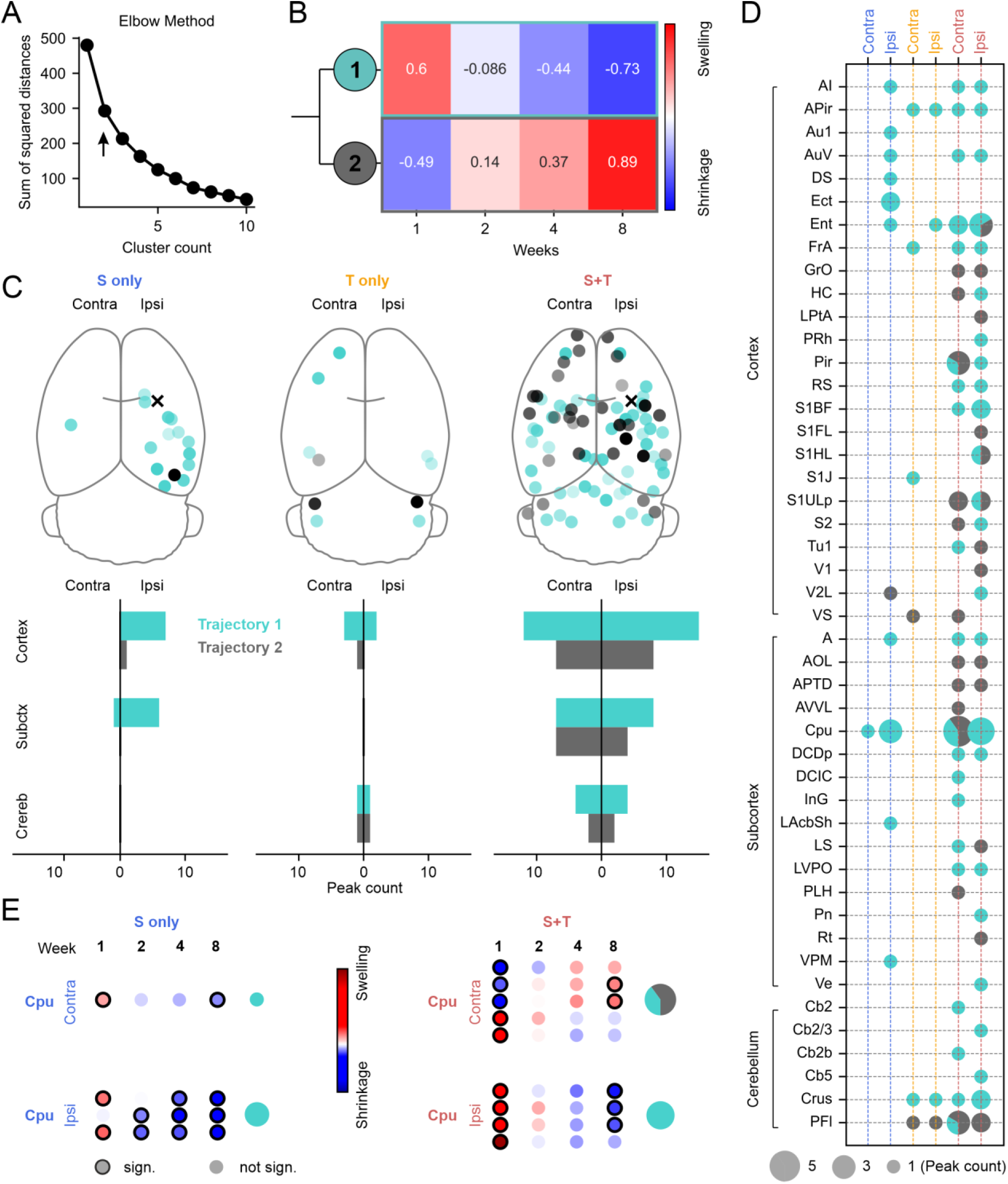
Volumetric changes occurring after stroke, training or the combination of both are characterized by two distinct temporal trajectories. (**A**) Elbow method identified two main clusters of volume changes over time. (**B**) Heat map displaying the two identified clusters (temporal trajectories) of volume changes over time. Numbers in squares indicate the strength of volume change ranging from -1 to 1 (**C**) Upper row: Three dimensional plot of the distribution of the two trajectories identified across the brain. Black crosses indicate stroke position. Lower row: Bar plots showing the count of peaks belonging to the first and second trajectory within the cortex, the subcortex and the cerebellum in both brain hemispheres. (**D**) Pie-charts displaying the percentage of each trajectory occurring in each brain region that displayed significant volume changes over time. For abbreviations see **Supplementary Table 1**. (**E**) Exemplary dot plots showing the trajectories of volume changes of all peaks detected within the Cpu. Values of percentage volume changes for each peak are provided in **Supplementary Table 2-4**. Data points with a black circle indicate significant swellings or shrinkages at the specific time point. Statistical analysis is described in the Methods (DBM).

In rats of all groups, we detected no obvious systematic spatial distribution of the different temporal trajectories (**Fig. 6 C**). In rats subjected solely to stroke, volume changes in both the cortex and subcortex primarily followed the first trajectory, with initial swellings transitioning to shrinkages (**Fig. 6 C**). Training alone induced volume changes that predominantly followed the first trajectory as well, especially in the cortex. However, in the cerebellum, both trajectories identified occurred in equal but limited proportions (**Fig. 6 C**). In rats that received both stroke and early multimodal training, the two trajectories were more evenly distributed across all the brain, although the majority of peaks still followed the first trajectory (**Fig. 6 C**). We did not find a bias of either temporal trajectory with respect to their distance from the stroke location (**Fig. S3**).

Importantly, post-stroke training prompted volume changes largely absent in the other groups: primarily, the second trajectory—initial shrinkage followed by subsequent swellings—was observed almost exclusively in rats of this group. Furthermore, several brain regions within the cortext, subcortex, and cerebellum exhibited volume changes belonging to both trajectories (**Fig. 6 D**), a pattern not observed in rats subjected only to stroke or motor training. For example, while all peaks in the ipsi- and contralateral Cpu of stroke-only rats followed the first trajectory, peaks in contralateral Cpu of post-stroke trained rats showed both trajectories (**Fig. 6 E**).

These data suggest that the distribution of volumetric temporal trajectories after stroke is determined by the type of intervention, with stroke or training alone typically inducing a pattern of initial swelling followed by shrinkage, whereas stroke with subsequent training leads to a more complex patterns where both swelling and shrinkage trajectories are observed across various brain regions.

## Discussion

In the present study, we found that early post-stroke multimodal motor training induces a recovery of task specific motor function and is associated with significant, brain-wide structural reorganization. Such changes were absent or only limited in rats subjected solely to stroke or motor training. Our data argue for profound effects of early multimodal motor training after stroke on both motor recovery and brain structural reorganization, highlighting the brain’s capacity to compensatorily and profoundly reshape during a limited time after stroke, if sufficiently stimulated (Witte, 1998).

Previous studies in rodents have suggested that environmental enrichment, which stimulates general bimanual limb use, combined with intensive task-specific motor training after motor stroke, improves both general and task-specific motor functions (Biernaskie et al., 2004; Jeffers and Corbett, 2018). However, environmental enrichment alone did not promote improvement of task-specific motor function (Jeffers and Corbett, 2018). In the present study, however, we demonstrate that multimodal motor training alone can cause motor improvements in a task-specific manner. Multimodal training on the parkour incorporated several therapeutic components with proven benefits in stroke recovery such as repetitive, temporally spaced and variable motor practice, increased difficulty and multisensory stimulation (Maier et al., 2019). Thus, we believe that multimodal motor training approaches not only improve performance in individual components of multimodal training, but also facilitate cognitive transfer to other, untrained motor tasks. This could imply that multimodal training enhances overall motor recovery and cognitive flexibility, promoting more comprehensive rehabilitation outcomes by leveraging the benefits of varied task engagement.

Volumetric brain changes in both rodents and humans have been found after motor training alone (Draganski et al., 2004; Sampaio-Baptista et al., 2013; Scholz et al., 2015b; Scholz et al., 2015a). In line with our results obtained in rats that exclusively underwent motor training, these studies found that volumetric alterations frequently occurred in a relatively low number of mostly task-specific brain regions. We detected volumetric alteration in brain regions such the cerebellar regions PFl and Crus. These regions are involved in synchronizing motor functions with sensory perception, vestibular reflexes (Beh et al., 2017), motor control and sensory motor behavior (Proville et al., 2014; Gaffield et al., 2022; Zhu et al., 2023) (**Fig. 3, 4**) and may hence, be relevant for trained task (parkour). Moreover, as our multimodal approach likely trains motor abilities symmetrically, we believe that this led to the observed bilateral volumetric changes in about 40% of the detected regions.

A few longitudinal studies in humans have suggested that stroke causes volumetric brain changes outside of the lesioned region (Brodtmann et al., 2012; Abela et al., 2015; Brodtmann et al., 2020; Bu et al., 2021; Syeda et al., 2022). However, many patients in these studies had already undergone some form of rehabilitation, making it challenging to attribute the observed volumetric effects solely to the stroke event itself. In experimental animals, it has been demonstrated that an isolated motor-stroke causes structural alterations in the cortex, hippocampus, and thalamus within the first 48 weeks post-stroke. However, region-specific analysis was not performed (Syeda et al., 2022). By performing atlas-based region-specific structural analysis, our study expands upon this previous knowledge. Although we did not observe structural changes in the hippocampus, our data are largely in line with Syeda et al., 2022, by showing changes in several brain regions located in the cortex and subcortex, including the thalamus (**Fig. 3, 4**). A notable brain region not detected by Syeda et al., 2022, but in our study, that showed bilateral volumetric changes in rats solely subjected to stroke is the caudate putamen (Cpu). This region is suggested to be involved in functions such as motor control and skill learning (Beaubaton et al., 1980; Yin et al., 2009; Yin, 2010; Kupferschmidt et al., 2017; Lemke et al., 2024). Interestingly, stroke-dependent loss of motor skills often forces patients to learn or acquire new skills as well, to compensate for the lost ones (Langhorne et al., 2009; Jones, 2017). Therefore, the structural changes observed within the Cpu of rats subjected solely to motor stroke may be associated with newly learned compensatory strategies, such as walking or sitting up to reach the drinking bottle or food within the home cage. Finally, similar to rats solely subjected to training the total number of regions with volume changes was relatively low, emphasizing that stroke itself only causes only limited volumetric changes across the brain.

In contrast to training or stroke alone, we found that the combination of both triggers substantial volumetric changes widely distributed across the entire brain (**Fig. 3, 4**). This indicates that stroke opens a window of increased potential for substantial structural reorganization, during which early multimodal motor training profoundly reshapes brain structure in a brain-wide manner. Volumetric alterations were detected within multiple cortical, subcortical, and cerebellar regions (**Fig. 3, 4**), with only a few of these regions showing changes in rats of the other groups. We believe it would be misleading to speculate why each of the regions and peaks that showed significant volumetric alterations may be important for motor recovery after stroke. Nevertheless, the abundance of changes detected is indicative of a unique post-stroke structural plasticity milieu (Murphy and Corbett, 2009; Zeiler and Krakauer, 2013; Jones, 2017; Campos et al., 2023) that strongly facilitates extensive structural reorganization on a brain-wide scale due to multimodal motor training. Conversely, multimodal motor training may also sculpt the post-stroke sensitive period plastic milieu (Zeiler and Krakauer, 2013).

What principles and mechanisms may underlie the structural changes observed? Generally, the root causes of volumetric structural brain changes, whether they involve swelling or shrinkage, remain enigmatic (Wenger et al., 2017). Moreover, increases in healthy brain volume have been temporally correlated with motor improvements (Draganski et al., 2004; Wenger et al., 2017), suggesting an association between volumetric changes and motor training. Motor training and other interventions often induce microstructural changes, such as synaptogenesis, changes in neuronal morphology, gliogenesis, glial volume changes, or vascular changes that often follow a similar time course as volumetric brain changes (Xu et al., 2009; Zatorre et al., 2012; Wenger et al., 2017; Schmidt et al., 2021; Sohn et al., 2022). Therefore, it can be assumed that structural brain changes observed in our interventional groups may be attributed to the same mechanisms that underlie brain plasticity after motor learning in the healthy brain.

Interestingly, especially in post-stroke trained rats, we found a surprisingly distinct temporal pattern: almost all brain regions that exhibited early swellings after stroke gradually shrank over the time-period examined (**Fig. 5, 6**). A part of the observed delayed shrinkage may be explained by the expansion-renormalization model which describes that a transient phase of structural swellings is followed by a return to baseline once a behaviorally optimal neuronal circuit has been selected (Wenger et al., 2017). However, our data show a further decrease in brain volume even below baseline levels in the following weeks after stroke (**Fig. 5, 6**). A potential explanation for this finding is that it may result from ongoing structural refinement within the circuits necessary for enhanced motor behavior.

While the aforementioned mechanisms may explain volume increases followed by decreases after stroke, literature offers fewer explanations for pure volume shrinkages after specific trainings, although they have been described repeatedly (Taubert et al., 2010; Takeuchi et al., 2011; Scholz et al., 2015a). In rats subjected to training alone, the general tendency for regional shrinkage, coupled with the rare occurrence of initial swellings, may be attributed to structural refinement of existing circuitry. The absence of difficulties in navigating the parkour suggests that these rats were not compelled to learn new motor skills, potentially explaining the infrequent initial swellings. Instead, it seems that these rats were only refining preexisting skills. In contrast, in post-stroke trained rats a significant portion of the two temporal trajectories was signified by early shrinkages followed by delayed swellings (**Fig. 5, 6**). The early shrinkages may be caused by structural refinement of existing circuitry and the delayed swellings may be a result of a later need of learning or re-learning motor skills as the difficulty of the parkour gradually increased. In other words, rats trained on the parkour post-stroke may experience two distinct motor learning periods - one early and one delayed. This explanation is supported by the finding that the time to finish the parkour tendentially increased with increasing difficulty (**Fig. 1**), suggesting the necessity of additional late motor adaptions due to increasing demand.

Why do all these widespread structural changes occur after post-stroke training? One possible explanation lies in the remapping hypothesis (Campos et al., 2023). This hypothesis suggests that spared circuits and regions remap over time to encode information previously encoded by those damaged due to stroke (Murphy and Corbett, 2009; Campos et al., 2023). Thus, the multi-regional changes observed after post-stroke training may reflect adaptation mechanisms within these regions, which consequently allow compensation or recovery of functions lost due to motor stroke. Moreover, as rats underwent multimodal motor training that encompassed multiple functions beyond pure motor skills, such as coordination, balance, and sensory-guided behavior, the plastic environment induced by stroke likely facilitated plastic changes in brain regions not directly involved in motor function. Alternatively, the abundance of volumetric changes found after stroke and early multimodal training may be explained by the redundancy or diaschisis theory. The redundancy hypothesis suggests that brain functions are duplicated or distributed across brain regions (Campos et al., 2023). Thus, the loss of a particular function after motor stroke is gradually overtaken by similar functions stored in the remaining brain regions. Diaschisis, on the other hand, describes a loss of long-range connectivity from the infarcted brain region to its projection areas (Buchkremer-Ratzmann et al., 1996; Campos et al., 2023). Stroke in the motor cortex would thus silence input in many brain regions typically innervated by the motor cortex. These regions would then plastically adapt to alterations in their input and may in turn undergo structural reorganization. The strong connectivity of the somatomotor cortex with a multitude of other brain areas, as revealed by neuronal tracing and functional investigations (Harris et al., 2019; Munoz-Castaneda et al., 2021; Weiler et al., 2024b; Weiler et al., 2024a), supports this hypothesis. In this context, we also want to emphasize the pronounced bilateral symmetry of structural alterations detected after post-stroke multimodal training. Given the tremendous connectivity between the two hemispheres (Gazzaniga, 2000; Munoz-Castaneda et al., 2021; Innocenti et al., 2022; Weiler et al., 2024a), stroke-induced plastic changes in ipsilateral brain regions may cause contralateral structural alterations in both homotopic and heterotopic areas as well. In reality, all these proposed mechanisms might work together, jointly contributing to both motor recovery and structural reorganization after a stroke (Campos et al., 2023). Notably, based on our data one could argue that multimodal motor training after stroke strongly facilitates such mechanisms, by effectively exploiting the plastic environment induced by stroke.

In conclusion, we here show that multimodal motor training initiated shortly after stroke significantly enhances the restoration of task-specific motor functions and causes extensive structural reorganization in a brain-wide manner. Hence, our findings highlight the importance of early, comprehensive and multimodal rehabilitation strategies in promoting brain plasticity and functional recovery post-stroke.

## Materials and Methods

### Animals

The study involved 12-week-old male albino Wistar rats (RccHan:WIST), housed in standard cages (2-5 animals) with a 12-hour light/dark cycle, *ad libitum* access to water and food at a controlled temperature of 21±1 °C and a humidity of 55±10%. Animal housing is regularly supervised by veterinaries for the state of Thuringia, Germany. All experimental procedures were conducted in accordance with the German Law on the Protection of Animals, the corresponding European Communities Council Directive (2010/63/EU) and approved by the Thüringer Landesamt für Lebensmittelsicherheit und Verbraucherschutz (Bad Langensalza, Germany). Importantly, every effort was made to minimize the number of animals and their suffering.

### Multimodal parkour training

Animals were trained on a parkour with a 10 cm wide runway, elevated 40 cm above the ground and equipped with 7 modifiable obstacles (teeterboard, ramp, alternating steps, palisade, fixed rods, suspension bridge, rotarod). These obstacles were used to enforce general limb use, balance, sensorimotor coordination and the execution of complex motor behaviors (multimodal training). Multimodal training was performed at 3 days per week, 5 times per day, with progressively increasing obstacle difficulty (**Fig. 1 A, B**). In weeks 0 and 1, animals were habituated to cross all obstacles at their easiest level. In weeks 2 to 4, the alternating steps, palisades, rods, suspension bridge and the rotarod were gradually modified towards their highest difficulty level to force gradual adaption to increasing demand. Modifications were introduced daily after the first run. In weeks 5 to 8, obstacles were randomly re-modified daily at a high difficulty level to further enforce gradual adaption. Importantly, the LR task, depicted in **Fig. 1 B**, at the end of the parkour was only installed when paw-placement behavior was assessed. Otherwise, during normal training, a normal jetty was inserted in its place.

#### Teeterboard

Generally, the board was tilted towards the start of the parkour (downwards). When crossed, it tilts towards the opposite side, which requires compensatory movement such as sudden balance adjustments.

#### Ramp

The ramp consisted of a shallow rise and steep descent. When crossing the top, animals were forced to move head-down. The tread surface consisted of a grid with distances greater than the tread of the animal feet, introducing a slipping risk.

#### Alternating steps

This obstacle consisted of 12 alternating steps with varying heights and distances. Increasing difficulty was achieved by decreasing the surfaces for paw placement, which simultaneously enforced an increase in the treat widths. Thereby, animals were challenged to adapt to unfamiliar limb positions to avoid falling.

#### Palisade

This obstacle consisted of six height-adjustable palisades. Difficulty was increased by introducing alternating heights of the palisades. This required the animals to climb up and down in narrow space while keeping correct paw placements to avoid falling.

#### Parallel rod

This obstacle consisted of two parallel round rods (diameter 5 mm) enforcing the animal to use the left paw for walking of the left and right paw for walking on the right rod. To increase difficulty, distance between the rods was increased and rods were loosened to allow for turning around their own longitudinal axis. Thus, animals were forced to use unfamiliar movement strategies while keeping the balance, to avoid falling.

#### Suspension bridge

This obstacle consisted of 8 steps each hanging on 4 chains. Initially, steps were connected by wires to fix them in their normal position (all steps on the same height-level). To increase difficulty we used 4 different strategies: 1) Wires connecting the steps were gradually removed to causing individual steps to swing. 2) By changing the length of the 4 chains height-differences between the steps were introduced. 3) Individual steps were replaced by steps smaller in size. 4) Steps were replaced by rods and two chains were removed resulting in the rods hanging on only two chains. Thus, this obstacle enforced the combination of several strategies, such as adapting the gait pattern to an unstable ground and keeping the balance to avoid falling.

#### Rotarod

This is a horizontal motorized rolling rod (diameter: 5cm). Initially, the rod did not turn around its own axis. To increase the difficulty to cross this rod, the rod was rolling around its own axis at increasing speed (1-5 rpm). Furthermore, the rolling direction was changed in a pseudorandom manner requiring animals to detect direction and speed of the rotation and to adapt their gait pattern, accordingly.

Importantly, for animals that felt down during the parkour, the timer was stopped and restarted after replacing them at the location they felt off. Generally, falling down happened very rarely in all groups of rats and was thus not further analyzed.

### Ladder rung (LR) walking task

Fore- and hind limb paw placement performance was analyzed utilizing the LR task (Metz and Whishaw, 2009). Baseline (BL) performance in all groups was assessed in week 0, one day before stroke induction in ‘S only’ and ‘S+T’ rats. Subsequently, rats of all groups were always tested on the 5^th^ day in weeks 1, 2, 4 and 8. On these days the task was optionally added at the end of the parkour. In rats which were not trained on the parkour we isolatedly assessed paw placement performance on the LR task. Rats of all groups were tested for 5 times on each testing day.

Animals crossed a 1-meter horizontal ladder consisting of clear Plexiglas side walls (19 cm high) and metal rungs (3 mm diameter) in variable distance (1 to 5 cm, **Fig. 1 B, 2A**). Locomotion was video-recorded with a high-speed camera (Pike F-32, Allied Vision Technologies, Germany). The camera was equipped with an 8-48 mm manual TV zoom lens (Pentax C60812, Pentax Rocoh Imaging Systems, Germany) and operated at 100 frames per second. Subsequently, paw placement performance was analyzed using a specific software (SIMI Motion, Simi Reality Motion Systems, Germany). Paw placement precision was rated using a 7-category quality score: (0) *Total miss*: deep fall after the limb completely missed a rung; (1) *Deep slip*: the limb slipped a rung and caused a fall; (2) *Slight slip*: the limb slipped a rung without interrupting the gait cycle; (3) *Replacement*: the limb was lifted to another rung before weight bearing; (4) *Correction*: the aimed rung or the limb position on the same rung was quickly corrected; (5) *Partial placement*: the limb was placed on a rung with either wrist/digits or heel/toes; (6) *Correct placement*: the mid-portion of a limb was placed on a rung with full weight support (Metz and Whishaw, 2009). Scores from the five trials were averaged and used for statistical analyses.

### Phototrombotic infarcation

Focal cortical lesions in rats solely subjected to stroke (‘S only’) and rats trained multi-modally after stroke (‘S+T’) were induced as described previously (Schmidt et al., 2012). Briefly, rats were anesthetized with isoflurane (2.0-2.5% in a 70:30 oxygen and nitrous oxide mixture), and a polyvinyl catheter was inserted into the tail vein. The skull skin was incised, and a fiber-optic bundle (Schott KL 1500, Germany, aperture: 2.5 mm) was positioned 0.5 mm anterior to bregma and 3.7 mm lateral to midline (**Fig. 1 E**) at the surface of the skull. This position is located above the right hemispheric somatomotor cortex covering parts of the somatosensory cortex processing information from the hind limb (S1HL) and fore limb (S1FL) and the motor cortex (primarily, the hind limb region). During the first minute of illumination (cold green light) rose Bengal (1.3 mg/100 g body weight, diluted in saline (0.9%), Aldrich Chemie, Steinheim, Germany) was injected. After 20 minutes of illumination, the catheter was removed, the skin was sutured, and animals were aerated with pure oxygen until they woke up. After 30 minutes of recovery in a warm environment, animals were returned to their home cages. Rats of the control group (‘Control’) and rats solely subjected to multimodal motor training (‘T only’) received the same anesthesia and surgery. However, instead of diluted rose Bengal pure saline was injected during the first minute of illumination (sham surgery). In rats of all groups the body temperature was controlled rectally and held constant at 37°C during the whole procedure.

### Magnetic resonance imaging (MRI)

To investigate whether rats from either group exhibited brain volumetric changes, we used MRI to repeatedly scan their brains. The first MRI, defined as the baseline (BL) measurement, was performed before any interventions and treatments began. Subsequent MRIs were conducted in weeks 1, 2, 4, and 8 (**Fig. 1 A**). The MRI procedures followed the protocol outlined by (Herrmann et al., 2012) and utilized a clinical whole-body scanner (Magnetom TIM Trio, Siemens Medical Solutions, Erlangen, Germany) equipped with a specialized rat head volume resonator featuring a linearly polarized Litz coil configuration (Doty Scientific Inc., Columbia, SC, USA). Rats were anesthetized with isoflurane (1.7% in oxygen, 1.5 l/min), and T_2_-weighted whole-brain images were acquired with a 3D SPACE sequence (Sampling Perfection with Application-Optimized Contrasts Using Different Flip Angle Evolutions, Siemens Healthcare, Erlangen, Germany) at 0.33 mm^3^ (voxel size) isotropic resolution. Key parameters were: matrix: 192 x 130 x 96 voxel, field of view: 64 x 43 x 32 mm, bandwidth 145 Hz/px, echo time (TE) 352 ms, repetition time (TR) 2500 ms, flip angle mode ‘T2var,’ echo spacing 10.7 ms, turbo factor 67, and partial Fourier 7/8 in both phase encode directions. The Doty coil improved signal-to-noise in comparison to standard Siemens coils and allowed a two-repetition protocol with a total acquisition time of 14 minutes.

### Deformation Based Morphometry (DBM)

Volumetric alterations of brain tissue were detected by using the DBM tool by (Gaser et al., 2012), integrated into MATLAB’s SPM8 (R2018a). This tool employs nonlinear registration by applying local deformations to rat brain images to align them (expansion or compression) with a reference image and encoding morphological disparities in 3D deformation fields. Deformation-specific displacement vectors (Jacobian determinants) were used to calculate volume differences at each voxel of the brain (**Fig. 1 D**). Sequential images of animal-individual data sets were rigidly registered to their BL image. Thereby, local deformations were introduced with high-dimensional nonlinear registration to identify subtle structural changes with great precision (Ashburner et al., 1999) (step1 in **Fig. 1 D**). To allow brain-region specific assignments of volumetric changes each imaged brain was aligned to the Paxinos rat brain Atlas (Paxinos and Watson, 2005) as described previously (Gaser et al., 2012). To finally assign each voxel to the coordinates of the Paxinos Atlas (Paxinos and Watson, 2005), the intra-pair deformations were nonlinearly transformed onto a customized and Paxinos-matched internal reference image by using the default spatial normalization method implemented in SPM8. Subsequently, displacement vectors were smoothed with a Gaussian kernel (FWHM of 0.8 mm) and used for whole-brain voxel-wise statistical analyses within the general linear model.

To detect individual voxels in the brains of each intervention group (‘S only’, T only’ and ‘S+T) that showed significant volumetric changes compared to the same localizations in the brains of control rats, we used F-contrasts to test for spatio-temporal interactions (F-test, RM ANOVA, threshold of p < 0.05, corrected for family-wise error [FWE]). This resulted in statistical maps revealing clusters corresponding to brain regions were the local structural changes (at least >10 voxels per cluster) significantly differed between the groups over time (step 2 in **Fig. 1 D,** and see also **Fig. 3 A**). In order to examine whether structural changes in each individual group were significant, and whether there were significant differences between the control and intervention groups (in weeks 1, 2, 4, and 8) we extracted the temporal trajectories cluster-peak-wise and analyzed them within and between groups using two-way RM ANOVA with Tukey HSD (step 3 in **Fig. 1D**). Importantly, for data presentation, and to define whether local volumes displayed shrinkage or swelling, all temporal trajectories of all interventional groups (for each animal) were corrected for the average control trajectories by subtraction. Thus, all brain region-based data depicted in **Fig. 3** and **4** were significant based on the abovementioned analysis. In **Fig. 5** and **6** we differentiated between significant and non-significant volumetric changes at each time point. However, all depicted temporal trajectories contained at least one time point were structural changes in a specific brain region reached significance.

### Estimation of stroke volume

Lesion size was assessed manually by a blinded investigator by the Cavalieri method, which enables precise and unbiased estimates of geometric quantities (Roberts et al., 2000). In brief, by using ImageJ, directly adjoining coronal brain sections (slice thickness: 0.33 mm) were overlaid by a regular square grid (grid space: 1.0 mm) and intersection points within and at the edges of the lesion were counted. Subsequently, the total volume of each lesion in all rats of the ‘S only’ and ‘S+T’ group was calculated. This analysis was performed in each week a MRI was performed. Data of individual rats in each group were averaged for data presentation.

### Statistical analysis

Details of all *n-*numbers are given in the first paragraph of the results and statistical analyses are provided in all figure captions and the Methods. The required sample sizes were estimated based on literature and our experience in performing similar experiments. Significance level was set as *p<*0.05 if not stated otherwise. Statistical analyses were performed in SPSS (IBM SPSS Statistics 29.0.0.0).

## Acknowledgements

Thanks are due to Ines Krumbein and Mike Hammer for excellent technical support and assistance. We thank Simon Weiler for helpful comments on our manuscript. The authors are further grateful to the support staff of the animal facility and of the Jena University Hospital. This work was supported by the European Union’s Horizon 2020 research and innovation program as part of the Marie Sklodowska-Curie grant agreement (SmartAge, grant 859890) to O.W.W., C.G. and C.F.; by the Carl Zeiss Foundation as part of the IMPULS project (grant P2019-01-006) to C.F. and C.G; by the German Research Foundation (DFG) as part of the Clinician Scientist Program OrganAge (WI 830/12-1 and WI 830/12-2, grant 413668513) to O.W.W., as part of the Excellence cluster “Precision medicine in chronic inflammation” (grant EXC2167) to C.F; by the Federal Ministry of Science and Education (BMBF) as part of the ERA PerMed (Pattern-Cog ERAPERMED2021-127) to C.G., as part of the Bernstein Focus (grant 01EO1002) to S.S., as part of the Jena Centre for Systems Biology of Aging (JenAge, grant 0315581) to C.F. and O.W.W; by the Interdisciplinary Center of Clinical Research of the Medical Faculty Jena (IZKF) with grant AMSP06 to A.U., grant MSP12 to S.G. and grant AMSP11 to M.T.; by the Foundation “Else Kröner-Fresenius-Stiftung“ within the Else Kröner Graduate School for Medical Students “Jena School for Ageing Medicine” (JSAM) and within the Else Kröner Research School for Physicians “AntiAge“ to O.W.W.

## Author contributions

SS, O.W.W., S.G and M.T. conceived the project. M.T. analyzed data, interpreted data, created Figures and wrote the manuscript. S.G. trained rats on the parkour, analyzed and interpreted data, and edited the manuscript. S.S. also trained rats on the parkour. K.H.H. and J.R.R. established the pipeline for MRI measurements. C.G. established the pipeline for DBM analysis. A.U., C.F. and K.H. interpreted data and edited the manuscript. S.S. and K.H.H. performed MRI measurements. S.S. and O.W.W. analyzed and interpreted data, edited the manuscript and supervised the project.

## Supplementary Information

**Supplementary Figure 1:**
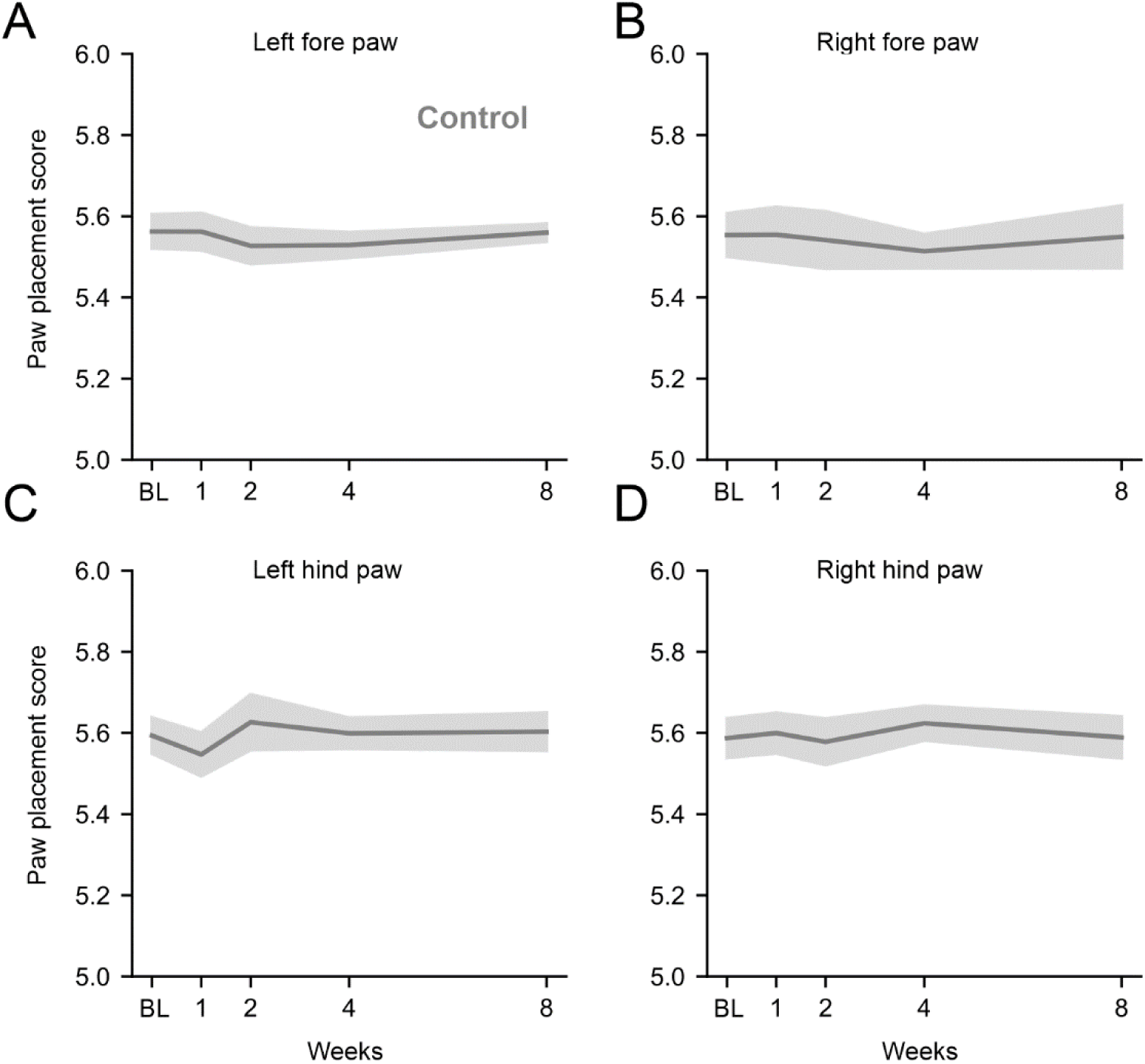
**Related to Fig. 2.** (**A**-**D**) Line plots showing the average paw placement score of control rats over time (± s.e.m.).

**Supplementary Figure 2:**
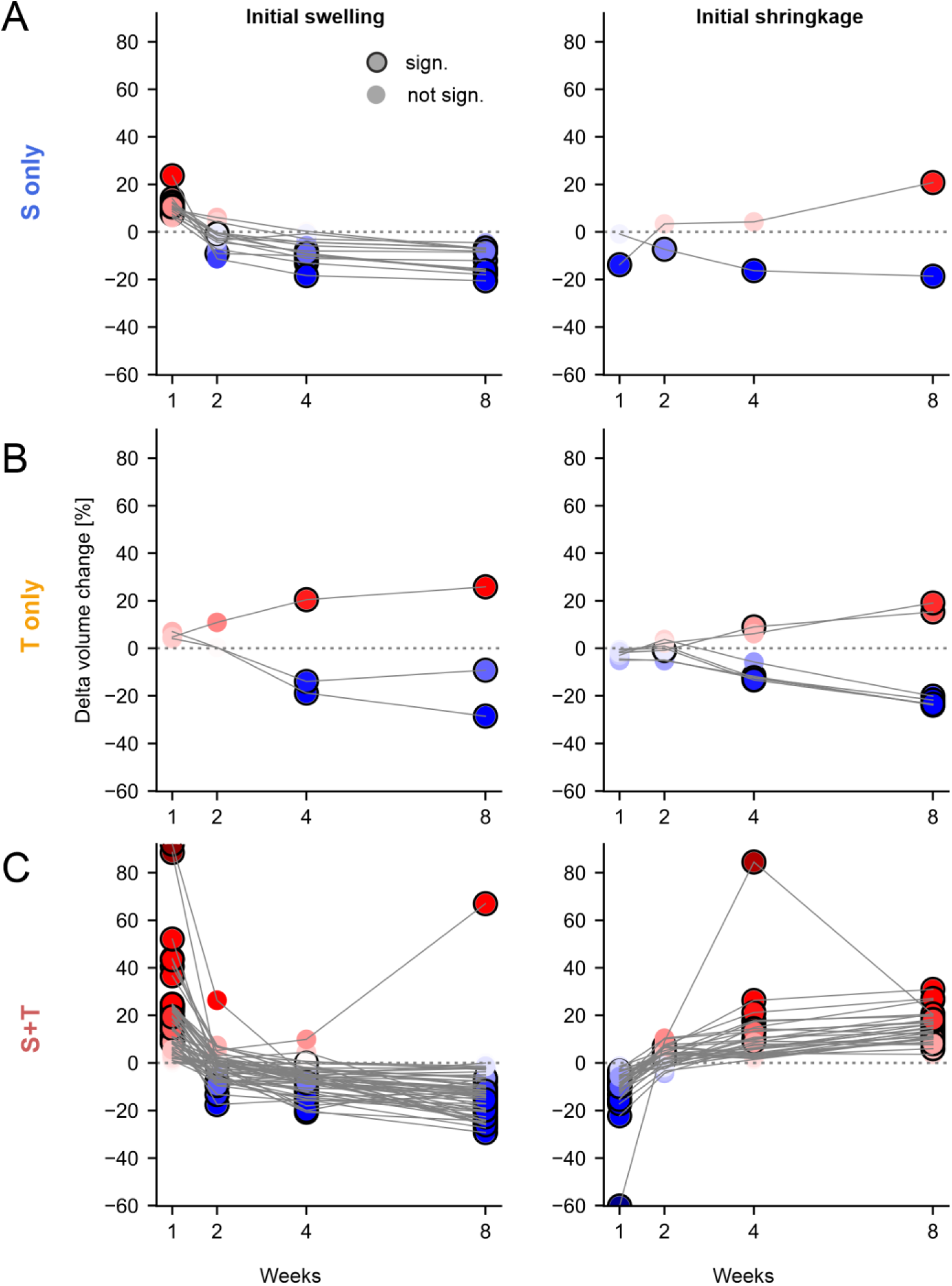
**Related to Fig. 5.** (**A**) Left: Line plot showing the volume changes over time of individual peaks that showed swellings at week one. Right: Line plot showing the volume changes over time of individual peaks that showed shrinkage at week one. Data points with a black circle indicate significant swellings or shrinkages at a specific time point (**B**, **C**) Same as in A. Values of percentage volume changes for each peak are provided in **Supplementary Table 2-4**.

**Supplementary Figure 3:**
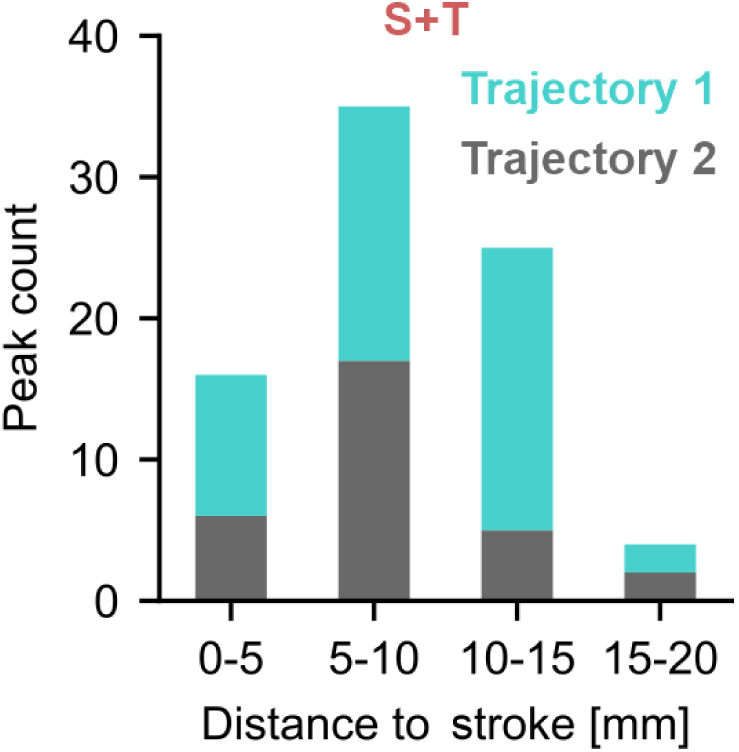
**Related to Fig. 6.** Bar plot showing the peak count for each temporal trajectory identified via k-means cluster analysis at different distances from the stroke.

**Supplementary Table 1:** Abbreviations of brain regions.

|  |  |
| --- | --- |
| **Cortex** |  |
| AI | Agranular insular cortex |
| APir | Amygdalopiriform transition area |
| Au1 | Primary auditory cortex |
| AuV | Secondary auditory cortex, ventral area |
| DS | Dorsal subiculum |
| Ect | Ectorhinal cortex |
| Ent | Entorhinal cortex |
| FrA | Frontal association cortex |
| GrO | Granular cell layer of the olfactory bulb |
| HC | Hippocampus |
| LPtA | Lateral parietal association cortex |
| Pir | Piriform cortex |
| PRh | Perirhinal cortex |
| RS | Retrosplenial cortex |
| S1BF | Primary somatosensory cortex, barrel field |
| S1FL | Primary somatosensory cortex, forelimb region |
| S1HL | Primary somatosensory cortex, hindlimb region |
| S1J | Primary somatosensory cortex, jaw region |
| S1ULp | Primary somatosensory cortex, upper lip region |
| S2 | Secondary somatosensory cortex |
| Tu1 | Olfactory tubercle |
| V1 | Primary visual cortex |
| V2L | Secondary visual cortex, lateral area |
| VS | Ventral subiculum |
| **Subcortex** |  |
| A | Amygdaloid area |
| AOL | Anterior olfactory nucleus, lateral part |
| APTD | Anterior pretectal nucleus, dorsal part |
| AVVL | Anteroventral thalamic nucleus, ventrolateral part |
| CPu | Caudate putamen (striatum) |
| DCDp | Dorsal cochlear nucleus, deep core |
| DCIC | Dorsal cortex of the inferior colliculus |
| InG | Intermediate gray layer of the superior colliculus |
| LAcSh | Lateral accumbens shell |
| LS | Lateral septal nucleus |
| LVPO | Lateroventral periolivary nucleus |
| PLH | Peduncular part of lateral hypothalamus |
| Pn | Pontine nuclei |
| Rt | Reticular thalamic nucleus |
| Ve | Vestibular nucleus |
| VPM | Ventral posteromedial thalamic nucleus |
| **Cerebellum** |  |
| Cb2b | 2b cerebellar lobule |
| Cb2 | 2nd cerebellar lobule |
| Cb2/3 | 2nd and 3rd cerebellar lobules |
| Cb5 | 5th cerebellar lobule |
| Crus | Crus of the ansiform lobule |
| PFI | Paraflocculus |
| **Ventricle** |  |
| 3V | 3rd ventricle |
| LV | Lateral ventricle |
| **White matter** |  |
| cc | Corpus callosum |
| cu | Cuneate fasciculus |
| fi | Fimbria of the hippocampus |
| ic | Internal capsule |
| m5 /s5 | Motor/sensory root of the trigeminal nerve |

**Supplementary Table 2:** Stroke only group. List of brain region related peaks that showed significant volumetric changes compared to the control group (F-test, RM ANOVA, threshold of *p-*values <0.05, corrected for family-wise error (FWE)). Values in the ‘Week’-columns indicate control corrected percentage volume changes compared to baseline (BL). Bold values indicate significance at the given time-points. Red colors indicate swellings and blue colors indicate shrinkages (*p*<0.05, peak-wise RM ANOVA with Tukey HSD).

| Brain region | Hemisph | Week 1 [%] | Week 2 [%] | Week 3 [%] | Week 4 [%] | F |
| --- | --- | --- | --- | --- | --- | --- |
| A | Ipsi | **9.4 ± 1.1** | **6.1 ± 1.3** | 0.3 ± 1.1 | **-7.1 ± 1.8** | 18.62 |
| AIP | Ipsi | **9.9 ± 1.2** | 4.2 ± 1.1 | -3 ± 1 | -4.6 ± 1.2 | 14.08 |
| Au1 | Ipsi | **7.4 ± 1.6** | -4.3 ± 1.7 | **-8 ± 1.2** | **-8.5 ± 1.5** | 15.05 |
| AuV | Ipsi | **9.4 ± 1.6** | -0.4 ± 1.1 | -4.4 ± 1.2 | -6.7 ± 1.2 | 15.24 |
| cc | Ipsi | **99.7 ± 14.9** | -4.2 ± 4.9 | **-9.8 ± 3.1** | -4 ± 6.4 | 28.61 |
| Cpu | Contra | **8.6 ± 1.2** | -2.6 ± 1.2 | -4.3 ± 1 | **-6.7 ± 0.9** | 20.58 |
| Cpu | Ipsi | **14 ± 2.8** | **-9.1 ± 1.3** | **-10.1 ± 1.4** | **-15.7 ± 1.2** | 22.43 |
| Cpu | Ipsi | -0.9 ± 1.9 | **-7.3 ± 1.3** | **-16.3 ± 1.6** | **-18.7 ± 2.2** | 15.87 |
| Cpu | Ipsi | **12.5 ± 2.5** | -0.4 ± 2 | **-9.4 ± 1.9** | **-16.7 ± 1.8** | 17.38 |
| DS | Ipsi | 6.1 ± 1.7 | -7.2 ± 1.8 | **-13.1 ± 2.2** | **-18 ± 2.2** | 16.87 |
| Ect | Ipsi | 6.6 ± 1.9 | -3.2 ± 1.4 | **-11.2 ± 1.7** | **-12.1 ± 1.5** | 19 |
| Ect | Ipsi | **12 ± 2.1** | -3.6 ± 1.3 | -0.8 ± 1.4 | -7.4 ± 1.7 | 15.86 |
| Ent | Ipsi | **11 ± 1.6** | -0.7 ± 1.4 | -5.8 ± 1.2 | **-8.1 ± 1.3** | 14.85 |
| ic | Ipsi | -8.4 ± 3 | 6.7 ± 3.4 | **19.7 ± 3.3** | **19.2 ± 3.1** | 13.90 |
| LAcSh | Ipsi | **10.5 ± 2.4** | -1.3 ± 1 | **-9 ± 1.8** | **-16.3 ± 1.7** | 16.49 |
| V2L | Ipsi | **-13.8 ± 1.4** | 3.4 ± 3.4 | 4.2 ± 2.4 | **20.8 ± 3.9** | 17.74 |
| VPM | Ipsi | **23.6 ± 5.1** | -11.3 ± 2.5 | **-18.4 ± 1.7** | **-20.5 ± 2.3** | 17.13 |

**Supplementary Table 3:**
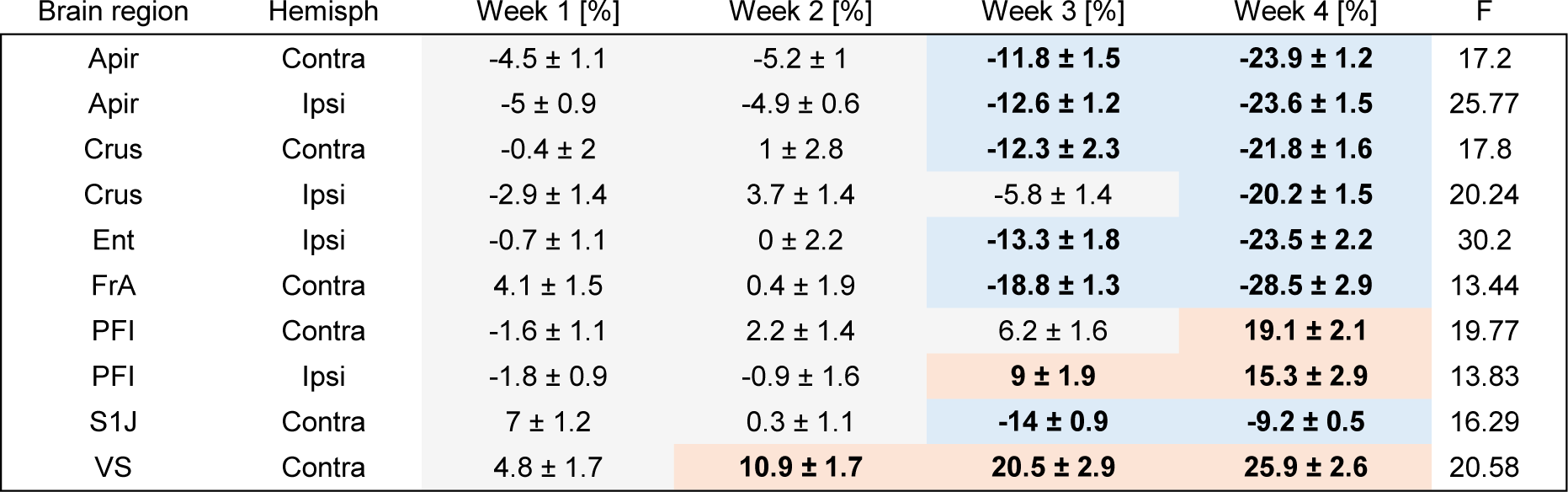
Training only group. List of brain region related peaks that showed significant volumetric changes compared to the control group (F-test, RM ANOVA, threshold of *p-*values <0.05, corrected for family-wise error (FWE)). Values in the ‘Week’-columns indicate control corrected percentage volume changes compared to baseline (BL). Bold values indicate significance at the given time-points. Red colors indicate swellings and blue colors indicate shrinkages (*p*<0.05, peak-wise RM ANOVA with Tukey HSD).

**Supplementary Table 4:** Stroke + Training group. List of brain region related peaks that showed significant volumetric changes compared to the control group (F-test, RM ANOVA, threshold of *p-*values <0.05, corrected for family-wise error (FWE)). Values in the ‘Week’-columns indicate control corrected percentage volume changes compared to baseline (BL). Bold values indicate significance at the given time-points. Red colors indicate swellings and blue colors indicate shrinkages (*p*<0.05, peak-wise RM ANOVA with Tukey HSD).

| Brain region | Hemisph | Week 1 [%] | Week 2 [%] | Week 3 [%] | Week 4 [%] | F |
| --- | --- | --- | --- | --- | --- | --- |
| 3V | Contra | **12 ± 1.5** | 0.6 ± 2.1 | -4.9 ± 3.2 | **-13.6 ± 1.8** | 19.55 |
| A | Contra | **13.6 ± 2.5** | 1.1 ± 0.7 | -3.9 ± 1.4 | -2.4 ± 1.1 | 17.4 |
| A | Ipsi | **9.1 ± 1** | 2.2 ± 0.9 | -3.7 ± 1.1 | **-7.9 ± 0.8** | 18.97 |
| AI | Contra | **10.7 ± 1.5** | 0.7 ± 0.6 | -0.2 ± 1.2 | -2 ± 1.5 | 14.82 |
| AI | Ipsi | **14.1 ± 4.4** | -7.2 ± 2.4 | -8.2 ± 2.1 | **-13.9 ± 2** | 14.53 |
| AOL | Contra | **-6.5 ± 0.9** | 4.3 ± 0.9 | 3.8 ± 1 | **8.9 ± 0.8** | 16.72 |
| AOL | Ipsi | **-13.2 ± 1.3** | **7.1 ± 1.2** | 4 ± 2.5 | 3.6 ± 2.3 | 20.21 |
| Apir | Contra | 3.9 ± 1.7 | -0.4 ± 1.2 | -5.4 ± 1.3 | **-14.7 ± 1.6** | 29.33 |
| Apir | Ipsi | 3.4 ± 2.6 | -7.2 ± 1.5 | **-15.4 ± 1.8** | **-27.3 ± 2.3** | 51.02 |
| APTD | Contra | -2.8 ± 1.8 | -2.6 ± 1.2 | **14.9 ± 2.4** | **26.5 ± 5.5** | 17.54 |
| APTD | Ipsi | -8.4 ± 2.3 | 4.1 ± 2.7 | **17.8 ± 3.2** | **19.7 ± 4.1** | 15.16 |
| AuV | Contra | **7.9 ± 3.1** | **-7.7 ± 1.3** | **-7.9 ± 1** | **-12.5 ± 1.5** | 17.44 |
| AuV | Ipsi | 1.1 ± 1.7 | **-7.7 ± 1.9** | **-15.2 ± 1.8** | **-20.5 ± 1.4** | 25.62 |
| AVVL | Contra | -8.2 ± 2.1 | 6.4 ± 4.8 | **26.3 ± 6.8** | **30.9 ± 6.2** | 17.72 |
| Cb2 | Contra | **23.5 ± 7.7** | -7.9 ± 3.7 | -5.3 ± 3.1 | **-24.1 ± 3** | 18.02 |
| Cb2/3 | Ipsi | **21.8 ± 4.3** | -4.6 ± 2.5 | -1.9 ± 2.6 | -9.4 ± 2.8 | 18.19 |
| Cb2b | Contra | **11 ± 2.2** | -6.8 ± 1.5 | **-10.5 ± 1.2** | **-12.7 ± 1.6** | 19.4 |
| Cb5 | Ipsi | **21.4 ± 2.4** | 1.1 ± 3.2 | -8 ± 2.2 | **-17 ± 2.1** | 28.73 |
| cc | Ipsi | **21.2 ± 4.1** | 1.3 ± 1.9 | **-8.6 ± 1.4** | -3.6 ± 2.6 | 23.49 |
| cc | Ipsi | **71.1 ± 9.7** | 3.3 ± 6.3 | -5.4 ± 3.7 | -9.3 ± 7 | 27.78 |
| Cpu | Contra | **23.6 ± 3.3** | 6.7 ± 1.4 | -2.2 ± 2 | -2.3 ± 1.8 | 29.09 |
| Cpu | Contra | **-22.2 ± 2.9** | -4.4 ± 1.6 | 7.9 ± 3.1 | 7.6 ± 3.6 | 16.53 |
| Cpu | Contra | **-15 ± 2.9** | 0.1 ± 1.5 | 10.7 ± 2.8 | **12.8 ± 3.4** | 16.13 |
| Cpu | Contra | **24.2 ± 6.2** | 1.2 ± 3 | -5.1 ± 1.6 | -3.7 ± 1.5 | 19.5 |
| Cpu | Contra | **-9.6 ± 1.3** | 2 ± 1.9 | 7.3 ± 2.3 | **10.9 ± 2.1** | 13.81 |
| Cpu | Ipsi | **24.3 ± 4.1** | -1.6 ± 2.5 | -8.4 ± 1.7 | **-13.7 ± 2.3** | 28.06 |
| Cpu | Ipsi | **43.9 ± 5.7** | 4.7 ± 2 | -6.1 ± 3.5 | **-10.3 ± 2.9** | 42.36 |
| Cpu | Ipsi | **40.5 ± 5.8** | 7.3 ± 4 | -5 ± 2.5 | **-11.3 ± 2** | 72.93 |
| Cpu | Ipsi | **88.5 ± 13.2** | -0.9 ± 4.6 | -6 ± 5.1 | -5 ± 2.6 | 60.45 |
| Crus | Contra | 0.1 ± 2.1 | -1.6 ± 1.4 | -0.1 ± 2.3 | **-24.3 ± 1.5** | 18.89 |
| Crus | Ipsi | 6.5 ± 2.7 | 0.9 ± 1.1 | -4.7 ± 1.7 | **-20.4 ± 2.2** | 23.03 |
| Crus | Ipsi | 0.1 ± 1.4 | 1.6 ± 1.3 | 4.5 ± 2 | **-19.4 ± 1.8** | 19.28 |
| cu | Ipsi | -6.5 ± 1.1 | -11 ± 1.6 | 6 ± 2.4 | **20.4 ± 2** | 18.96 |
| DCDp | Contra | **10.8 ± 2.9** | -4.7 ± 2.3 | -3.9 ± 1.7 | **-25.8 ± 1.7** | 28.92 |
| DCDp | Ipsi | 9.1 ± 2.5 | -5.2 ± 1.6 | -7.1 ± 1.4 | **-25.3 ± 1.5** | 25.9 |
| DCIC | Contra | 8.2 ± 3.9 | -0.4 ± 3.4 | -6.2 ± 3.3 | **-25.3 ± 2.6** | 21.3 |
| Ent | Contra | 3 ± 1.6 | -0.2 ± 1.4 | **-6.9 ± 1.3** | **-11.7 ± 2.2** | 14.61 |
| Ent | Contra | 1.8 ± 2.4 | -0.5 ± 2.8 | -8.6 ± 2 | **-20.7 ± 2** | 23.18 |
| Ent | Ipsi | 1 ± 1 | -2.8 ± 1.5 | **-12.5 ± 1.4** | **-23.8 ± 2** | 35.2 |
| Ent | Ipsi | **23.1 ± 2.7** | 2 ± 1.7 | -5.9 ± 1.8 | **-11.4 ± 1.6** | 29.58 |
| Ent | Ipsi | -3.5 ± 1.1 | 3.8 ± 1 | 1.8 ± 0.8 | **10.7 ± 1.4** | 20.12 |
| fi | Contra | 4 ± 2.4 | **-9.8 ± 1.5** | **-18.4 ± 1.3** | **-18.8 ± 1.4** | 21.97 |
| FrA | Contra | 2.2 ± 1.9 | -0.1 ± 2 | **-20 ± 1.6** | **-29.3 ± 3.1** | 14.31 |
| FrA | Ipsi | 6.1 ± 3.1 | -4.2 ± 3 | **-20.7 ± 2.2** | **-24.4 ± 2.8** | 20.1 |
| GrO | Contra | -3.6 ± 1.5 | 1.4 ± 2.8 | 6.3 ± 2.3 | **14.6 ± 2.7** | 17.74 |
| GrO | Ipsi | **-9.3 ± 2** | 4.1 ± 3.6 | 6 ± 3.5 | **16.3 ± 3.9** | 21.44 |
| HC | Contra | 8.5 ± 5.9 | -8.4 ± 1.8 | -6.5 ± 2.9 | **-26 ± 1.7** | 15.84 |
| HC | Ipsi | **92 ± 14.1** | **26.3 ± 5.9** | -5.8 ± 3.3 | 0 ± 3.6 | 47.71 |
| InG | Contra | **22.6 ± 5.1** | -1.4 ± 1.7 | -3.9 ± 1.7 | -2.3 ± 2.1 | 24.99 |
| LPtA | Ipsi | **-10.9 ± 1.5** | 1.6 ± 2.5 | **17.3 ± 4.1** | **20.4 ± 4.9** | 25.8 |
| LS | Contra | **24.4 ± 5.2** | 2.7 ± 1.9 | -7.2 ± 2.4 | -10.4 ± 2 | 17.48 |
| LS | Ipsi | **-10.5 ± 3.1** | 7.6 ± 2.6 | **21.2 ± 3.3** | **27.1 ± 5.2** | 20.59 |
| LV | Contra | **43.2 ± 10.5** | 2.4 ± 3.6 | -6.9 ± 1.7 | -8 ± 1.5 | 28.4 |
| LV | Ipsi | **28.8 ± 6.5** | -5.6 ± 2.7 | **-11 ± 1.7** | **-23.1 ± 2.9** | 32.39 |
| LV | Ipsi | **139.6 ± 15.5** | **20.9 ± 7.1** | -16.3 ± 4.5 | -14.7 ± 6.2 | 100.6 |
| LV | Ipsi | **35.8 ± 2.7** | **11.2 ± 3.2** | 2.1 ± 2.7 | 1.4 ± 1.5 | 47.95 |
| LVPO | Contra | **19.9 ± 4.7** | -2.6 ± 3.5 | -8.5 ± 2.8 | -13.4 ± 3.8 | 15.19 |
| LVPO | Ipsi | **19 ± 4.2** | -9.5 ± 3.3 | **-14.2 ± 2.1** | **-12.5 ± 3** | 21.06 |
| m5 > c5 | Contra | **18 ± 3.9** | 0 ± 2.1 | -8.6 ± 2.1 | **-12.7 ± 1.9** | 20.51 |
| PFI | Contra | -5.3 ± 1.1 | -1.6 ± 1.7 | 3.6 ± 1.2 | **13.7 ± 1.4** | 16.75 |
| PFI | Contra | -5.9 ± 1 | -0.1 ± 1.6 | 5.6 ± 1.8 | **15 ± 2.3** | 16.64 |
| PFI | Contra | 5.4 ± 2.3 | -6.7 ± 1.7 | **-11.4 ± 2.2** | **-22.7 ± 1.7** | 15.32 |
| PFI | Ipsi | -1.4 ± 1.1 | 5.1 ± 1 | **13.5 ± 1.2** | **16.9 ± 1.5** | 21.21 |
| PFI | Ipsi | -1.5 ± 1.7 | -0.5 ± 1.8 | **10.2 ± 2** | **16.1 ± 3.4** | 16.52 |
| Pir | Contra | **-12.1 ± 1.1** | 1.5 ± 1.2 | 2.6 ± 1 | **12.7 ± 2.8** | 13.88 |
| Pir | Contra | **24.7 ± 4.4** | 3.4 ± 2.9 | 0.7 ± 1.8 | -2.9 ± 1.1 | 24.75 |
| Pir | Contra | -7 ± 1.4 | 6.4 ± 1.9 | 4.1 ± 1.8 | **13.4 ± 3** | 15.44 |
| Pir | Ipsi | **19.2 ± 3.4** | -2.9 ± 1.7 | -2.8 ± 2.4 | -5.9 ± 1.4 | 24.17 |
| PLH | Contra | **-9.3 ± 2.6** | **10.4 ± 1.8** | **14.4 ± 2.4** | **12 ± 2.2** | 24.05 |
| Pn | Ipsi | **36.6 ± 6.3** | 6.7 ± 4.8 | -2 ± 4.1 | -10 ± 2.4 | 23.26 |
| PRh | Ipsi | 4.6 ± 3 | -3.3 ± 1.6 | **-11.7 ± 1.8** | **-19 ± 1.5** | 28.6 |
| RS | Contra | **17.3 ± 5.1** | -2.1 ± 1.9 | **-18.1 ± 2.8** | **-12.5 ± 3.3** | 19.24 |
| RS | Ipsi | **43.4 ± 8.2** | 0.2 ± 1.9 | -5.8 ± 1.6 | -7.5 ± 2.8 | 29.75 |
| Rt | Ipsi | 11 ± 3.5 | 5.2 ± 3.2 | 9.9 ± 6.7 | **67 ± 16.2** | 23.44 |
| S1BF | Contra | **8.5 ± 1.5** | 0.6 ± 0.7 | -2.4 ± 1.2 | **-6.4 ± 1.4** | 17.82 |
| S1BF | Ipsi | 8.9 ± 6.6 | **-13.3 ± 2.5** | **-15.9 ± 2.6** | **-14.4 ± 2.2** | 15.41 |
| S1BF | Ipsi | **15.8 ± 4.8** | **-17.5 ± 1.8** | **-15.4 ± 2.3** | **-17.2 ± 2.2** | 47.98 |
| S1FL | Ipsi | **-17.1 ± 1.2** | -3.4 ± 1.6 | 1.4 ± 2.5 | **10.5 ± 3.3** | 20.21 |
| S1HL | Ipsi | **-60.2 ± 3.8** | 9.1 ± 2.3 | **84.5 ± 8.1** | 20.3 ± 6.2 | 86.17 |
| S1HL | Ipsi | **52.1 ± 8.7** | -5.2 ± 3.4 | **-20 ± 6.9** | -6.4 ± 7.9 | 22.03 |
| S1ULp | Contra | **-5.1 ± 1.6** | 3.5 ± 2 | 3.3 ± 1.2 | **10 ± 2** | 24.18 |
| S1ULp | Contra | -3.5 ± 0.7 | 4.1 ± 2.1 | 2.2 ± 0.9 | **10.7 ± 1.8** | 16.91 |
| S1ULp | Ipsi | **14.5 ± 5.4** | 0.1 ± 4.5 | -8.9 ± 3.9 | **-11.9 ± 2.9** | 18.21 |
| S1ULp | Ipsi | **-11.5 ± 2.4** | -1.8 ± 3.1 | 5.8 ± 4.2 | **11.3 ± 3.2** | 13.45 |
| S2 | Contra | **-9.3 ± 0.8** | 3.8 ± 1.5 | 3.1 ± 1 | **6 ± 0.8** | 34.48 |
| S2 | Ipsi | 3.2 ± 3.5 | -5.9 ± 2.3 | **-14.4 ± 1.9** | **-19.8 ± 1.5** | 23.27 |
| Tu1 | Contra | **19.3 ± 3.5** | 6.8 ± 1.5 | 0.4 ± 1.2 | -1.4 ± 1.2 | 19.07 |
| Tu1 | Ipsi | **-15.3 ± 2.3** | 7.6 ± 4.3 | 9.7 ± 3.9 | **20.6 ± 4.4** | 16.82 |
| V1 | Ipsi | **-9.9 ± 1.7** | 6.4 ± 1.5 | **9.2 ± 1.6** | **8.1 ± 1.6** | 19.13 |
| V2L | Ipsi | 5.5 ± 1.4 | **-9.5 ± 1.6** | **-8.2 ± 1.9** | **-15.5 ± 0.7** | 17.16 |
| Ve | Ipsi | -5.8 ± 1.4 | -1.4 ± 1.8 | 8.6 ± 2.2 | **18.3 ± 2.7** | 18.08 |
| VS | Contra | -2.5 ± 1.2 | **10.1 ± 1.3** | **13.1 ± 1.7** | **17.4 ± 2.1** | 21.93 |

